# Unbalancing cAMP and Ras/MAPK pathways as a therapeutic strategy for cutaneous neurofibromas

**DOI:** 10.1101/2022.12.23.521754

**Authors:** Helena Mazuelas, Míriam Magallón-Lorenz, Itziar Uriarte-Arrázola, Alejandro Negro, Inma Rosas, Ignacio Blanco, Elisabeth Castellanos, Conxi Lázaro, Bernat Gel, Meritxell Carrió, Eduard Serra

**Affiliations:** Hereditary Cancer Group, Germans Trias i Pujol Research Institute (IGTP); Can Ruti Campus, Badalona, Barcelona, 08916; Spain; Clinical Genomics Research Group, Germans Trias i Pujol Research Institute (IGTP); Can Ruti Campus, Badalona, Barcelona, 08098; Spain; Genetics Service, Germans Trias i Pujol University Hospital (HUGTiP), Can Ruti Campus, Badalona, Barcelona, 08098; Spain; Hereditary Cancer Program, Catalan Institute of Oncology (ICO-IDIBELL), L’Hospitalet de Llobregat, Barcelona, 08098; Spain; Centro de Investigación Biomédica en Red de Cáncer (CIBERONC), Spain; Departament de Fonaments Clínics, Facultat de Medicina i Ciències de la Salut, Universitat de Barcelona (UB), 08036, Barcelona, Spain

## Abstract

Cutaneous neurofibromas (cNFs) are benign Schwann cell (SC) tumors arising from subepidermal glia. Neurofibromatosis Type 1 (NF1) individuals may develop thousands of cNFs, greatly affecting their quality of life. cNF growth is governed by the proliferation of *NF1*(-/-) SCs, highly influenced by the interaction with a *NF1*(+/-) microenvironment, consisting of fibroblasts (FBs), immune cells, etc. To decompose crosstalk between SCs and the microenvironment we used single cultures and co-cultures of cNF-derived SCs and FBs and identified an expression signature specific to SC-FB interaction. This signature was enriched in genes involved in immune cell migration, that were functionally validated by secretion analysis of SC-FB co-cultures, suggesting a role of SC-FB crosstalk in immune cell recruitment. The signature also captured components of different developmental signaling pathways, among them, the cAMP elevator G protein-coupled receptor 68 (*GPR68*). Activation of Gpr68 by Ogerin reduced the viability and proliferation of cNF-derived SCs and SC-FB co-cultures. Moreover, Ogerin in combination with the MEKi Selumetinib induced loss of viability, SC differentiation, and death. These results were corroborated using an iPSC-derived 3D neurofibromasphere model. The unbalancing of the Ras and cAMP pathways by combining a MEKi and a cAMP elevator arises as a potential treatment for cNFs.

## Introduction

Cutaneous neurofibromas (cNFs) are benign Schwann cell (SC) tumors that originate in the peripheral nervous system that resides in the skin, most likely in the sub-epidermal glia (Radomska et al., 2019). They form discrete nodules that can adopt different forms depending on distinct variables: the exact location they arise within the skin; the skin type of the patient; the age of onset; the growth rate; etc. (Ortonne et al., 2018). cNFs normally appear during puberty and are present in more than 95% of Neurofibromatosis Type 1 patients (Ortonne et al., 2018). Their number is greatly variable depending on the NF1 individual, ranging from tens to thousands, but always increasing throughout life (Cannon et al., 2018; Duong et al., 2011; Huson et al., 1988). cNFs might be itchy or painful, but most are asymptomatic. Although cNFs do not progress to malignancy and rarely are associated with clinical complications, they can greatly impact the quality of life of NF1 patients due to disfigurement, dysesthesia, and the psychological impact of the perceived disease visibility (Granstörm et al., 2012; Wolkenstein et al., 2001). Current clinical management of cNFs involves the surveillance of lesions and may require treatment by surgical resection, laser treatment, electrodesiccation, or other means (Peltonen et al., 2022; Verma et al., 2018). The development of effective drugs for treating cNFs is currently an active research area in the field. The MEKi Selumetinib is being assayed systemically and topically (de Blank et al., 2022). However, further development of new effective drugs and delivery methods is highly needed.

A typical histological description of a cNF is that of a tumor with abundant mucoid stroma, floating strands of collagen, and scarce cells (Stemmer-Rachamimov and Nielsen, 2012). Despite hypocellularity, cNFs are composed of multiple cell types. Typically, SCs comprise between 40% and 80% of all cellular content (Krone et al., 1983; Peltonen et al., 1988). The next big cell component is fibroblast (FB) from the endoneurium. cNFs also contain a rich immune infiltrate, composed of mast cells (Kallionpää et al., 2021), lymphocytes, macrophages, and others. Endothelial cells, perineural cells, and axons are also present, all embedded in a collagen-rich extracellular matrix (Konomi et al., 1989; Uitto et al., 1986). Among the different cell types composing cNFs, only SCs bear a double inactivation of the *NF1* gene (Maertens et al., 2006; Serra et al., 2000)making them susceptible to over-proliferation. However, much evidence obtained *in vitro* and using genetically modified animal models draw an essential role of the rich *NF1*(+/-) microenvironment in cNF (revised in (Jiang et al., 2021). Whether the *NF1*(+/-) status of the microenvironment is essential or just a facilitator of cNF development is still under debate since different mouse models bring distinct responses. In humans, sporadic plexiform neurofibromas carrying a biallelic inactivation of *NF1* have been reported in individuals with the rest of the cells being *NF1* WT cells (Beert et al., 2012). Anyhow, in the NF1 context, the microenvironment plays, at least, a facilitator role in cNF growth. The most studied component of the microenvironment has been the immune infiltrate, and its role in promoting neurofibroma growth (revised in (Fletcher et al., 2020)), in which different immune cells, like mast cells, macrophages, T-cells, etc., have been involved in one way or another. The complex interactions that might exist between SCs and the immune system, but also with the rest of cell components (FBs, axons) need to be further experimentally dissected and clarified. One way of studying cell-cell interactions is the use of cNF-derived primary cell cultures, comparing their biology either in single cultures or in co-cultures. cNF cell components can be dissociated and either sorted by flow cytometry or panning strategies (Choi et al., 2017), or by differential growing conditions in culture. In this regard, conditions for selectively expanding *NF1*(-/-) SCs and *NF1*(+/-) FBs from cNFs are well established (Serra et al., 2000).

Neurofibromin’s role in Ras/MAPK regulation is well characterized (Ratner and Miller, 2015), also in the context of SCs (Kim et al., 1995; Sherman et al., 2000). The role of neurofibromin in the cAMP pathway has also been determined (The, 1997; Tong et al., 2002) although much less understood at the molecular level. Recently, cAMP intracellular elevation has been involved in the reduction of SC precursor self-renewal and cNF SC proliferation (Patritti-Cram et al., 2022). In agreement, *NF1*(+/-) and *NF1*(-/-) cNF-derived SC cultures can be obtained basically by modifying the exposure to cAMP-elevating agents, since *NF1*(-/-) SCs decrease proliferation after long expositions to elevated cAMP compared to *NF1*(+/-) SCs (Serra et al., 2000). This could be somehow related to the increased basal cAMP levels in *Nf1*-deficient mouse SCs compared to *Nf1*(+/-) (Kim et al., 2001). In addition, the level of cAMP signaling is crucial for switching from a proliferative state, to a myelin-forming differentiation state (Arthur-Farraj et al., 2011). Both the Ras/MAPK and the cAMP/PKA pathways need to be active for prolific SC proliferation (Rahmatullah et al., 1998).

In the present work, we have investigated the crosstalk between cNF-derived *NF1*(-/-) SCs and *NF1*(+/-) FBs. We identified a transcriptional signature due to the SC-FB interaction, enriched in genes involved in immune cell migration and developmental signaling pathways. The identification of the cAMP elevator G protein-coupled receptor 68 (*GPR68*) prompted us to test the effect of elevating cAMP in *NF1*(-/-) SCs, alone or in combination with the MEKi Selumetinib. Results point to the inhibition of the Ras pathway and activation of the cAMP pathway as a potential therapeutic strategy for cNFs.

## Results

We developed an experimental framework to investigate cNF-derived SC-FB crosstalk, based on the identification of transcriptional changes in both cell types specifically produced by their interaction (**Figure 1A**). We first identified both constitutional and somatic *NF1* mutations present in cNFs (**Table 1**) and set up experimental conditions to analyze the transcriptome of the SC-FB interaction.

**Figure 1.**
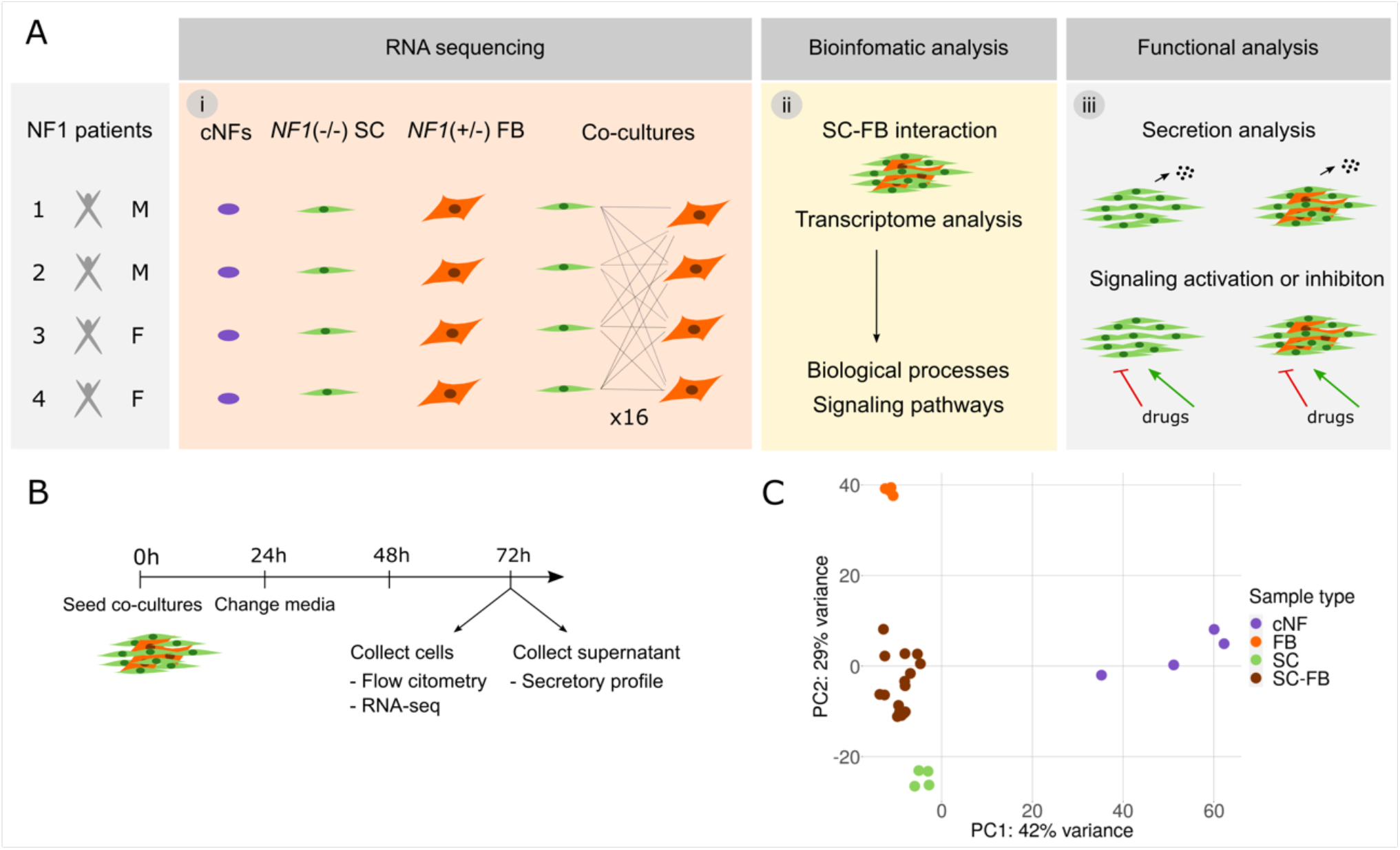
Experimental set up workflow. A. Schematic representation showing the three different phases of the project: (i) establishment of SC and FB cultures from cNFs and establishment of SC-FB co-cultures and RNA-seq,(ii) bioinformatic analysis, and (iii) functional analysis. B. Timeline representation of the co-culture experiment. C. Principal component analysis of cNFs, SCs and FBs single cultures, and SC-FB co-cultures from RNA-seq data showing a clear separation of cell types. SC: Schwann Cells; FB: Fibroblasts; SC-FB: SC-FB co-cultures, cNF: cutaneous neurofibroma.

**Table 1.** Summary of constitutional and somatic *NF1* alterations in selected cNFs

| *Patient information* |  |  | *Tumor information* |  |  |
| --- | --- | --- | --- | --- | --- |
| Patient ID | Sex | *NF1* constitutional mutation | Tumor ID | Age at resection | *NF1* somatic mutation |
| 1 | M | c.5898_5899delGA | cNF1_1 | 46 | c.4120C>T |
| 17 | M | c.3826C>T | cNF1_2 | 34 | c.4537C>T |
| 27 | F | c.4309G>T | cNF1_3 | 34 | c.810_833delAATCATTCTCCTTATCTTGTGTCC |
| 28 | F | c.3233C>G | cNF1_4 | 76 | c.3158C>G |

### Transcriptome analysis of cNFs and cNF-derived cell cultures and co-cultures

We obtained cNFs from 4 independent NF1 individuals, two females and two males (**Table 1**, and M&M section) and derived single cultures of *NF1*(-/-) SCs and *NF1*(+/-) FBs of each tumor (Serra et al. 2000). We sequenced the entire *NF1* coding region (Castellanos et al. 2017) from these cultures and demonstrated their constitutional or somatic nature by comparing *NF1* specific mutations also in *NF1*(+/-) FB cultures. We also applied microsatellite analysis for LOH identification in the *NF1* region (Garcia-Linares et al. 2012). Among the distinct cNFs analyzed, we selected 4 cNFs from 4 different NF1 patients (**Table 1**), performed whole exome sequencing of both cNF-derived SC and FB cultures, and analyzed all constitutional genomic variance present in coding genes as well as all somatic mutations present in *NF1*(-/-) SCs. As previously reported (Ferrer et al., 2018), we did not identify any significant recurrently mutated gene present in NF1-associated cNF-derived SCs, besides the *NF1* gene.

Using single cultures of SCs and FBs derived from all cNFs, we set up conditions for generating SC-FB co-cultures with a final defined proportion of cells resembling those present on average in neurofibromas, approximately 60-70% SCs and 30-40% FBs **(Supplemental Figure 1;** Carrio et al. 2018). For co-culture experiments, we controlled cell density over time and assessed the presence of both cell types using different markers. To capture biological differences coming from the distinct cNFs, we generated 16 SC-FB co-cultures by mixing each SC culture with FBs derived from each of the 4 cNFs (**Figure 1A**). Both single cultures and co-cultures were submitted to experimental conditions (**Figure 1B**), and after 72h, cells and supernatants were collected. We used part of the harvested cells to quantify the percentage of both cell types, through flow cytometry analysis using the SC maker p75 (*NGFR*), thus keeping track of the exact proportions in single cultures and co-cultures (**Supplemental Figure 2**). We extracted RNA and performed RNA-seq for all 16 SC-FB co-cultures, the 4 single SC cultures, the 4 single FB cultures, and the 4 original cNFs. Principal component analysis (PCA, **Figure 1C**) nicely grouped samples of each type and separated all different groups of samples. The two main components capturing most parts of the variability among samples evidenced similarities and differences between SC-FB co-cultures and single cultures, as well as between SC-FB co-cultures and cNFs.

### An expression profile of SC-FB crosstalk

From all genes expressed in SC-FB co-cultures, we aimed to identify a transcriptional signature specifically representing genes whose expression was dominated by the interaction between SCs and FBs. For that, we set up a specific bioinformatic analysis (**Figure 2A**). For each of the 16 SC-FB co-cultures, a virtual companion co-culture was generated by the random sampling of reads from the two SC and FB single cultures used for each co-culture generation. For instance, for the co-culture SC1-FB3, the random sampling of reads was performed from RNA-seq data of single SC1 and FB3 cultures (**Supplemental Figure 2**). The proportion of reads sampled from every single culture was equivalent to the exact proportion of SCs and FBs present in each co-culture, determined by flow cytometry analysis. In this regard, companion virtual co-cultures could be seen as co-cultures with no expression changes due to SC-FB interactions, containing the same expression in SCs and FBs as in their respective single cultures although in a mixed composition. PCA revealed a high expression similarity between real and virtual co-cultures (**Figure 2B**) although it also left a margin for consistent expression differences. To identify an expression profile specific to SC-FB crosstalk, a differential expression analysis between real and virtual co-cultures was performed and results were plotted in a heatmap representing an unsupervised cluster analysis of differentially expressed genes (**Figure 2C**). This analysis identified a transcriptional signature containing upregulated and downregulated genes in real vs virtual co-cultures (**Supplemental File 1**). To validate this expression profile in a different neurofibroma model system, we selected all upregulated genes and analyzed their expression in an iPSC-derived 3D neurofibroma model, consisting of spheroids containing only SCs or containing an admixture of SCs and FBs (Mazuelas et al. 2022). An unsupervised cluster analysis of this upregulated gene signature in real and virtual co-cultures, SCs and SC-FB spheroids, and cNFs, indicated that most genes of the signature were also specifically expressed in SC-FB spheroids, clustering together with real SC-FB co-cultures (**Supplemental Figure 3**). In comparison, only a small part of the genes was expressed in SC-only-spheroids, which clustered together with virtual co-cultures (**Supplemental Figure 3**). Thus, we identified a robust SC-FB crosstalk signature that was preserved in different neurofibroma co-culture systems. We next aimed to translate the identified SC-FB crosstalk signature into biological processes and signaling pathways. For that, we selected all genes constituting the signature and performed an enrichment analysis (**Figure 3, Supplemental File 2**). We selected two groups of biological processes: 1) those related to immune response, immune cell migration, and chemotaxis; 2) those broadly grouped as commonly involved in developmental processes and developmental signaling pathways (**Figure 3**).

**Figure 2.**
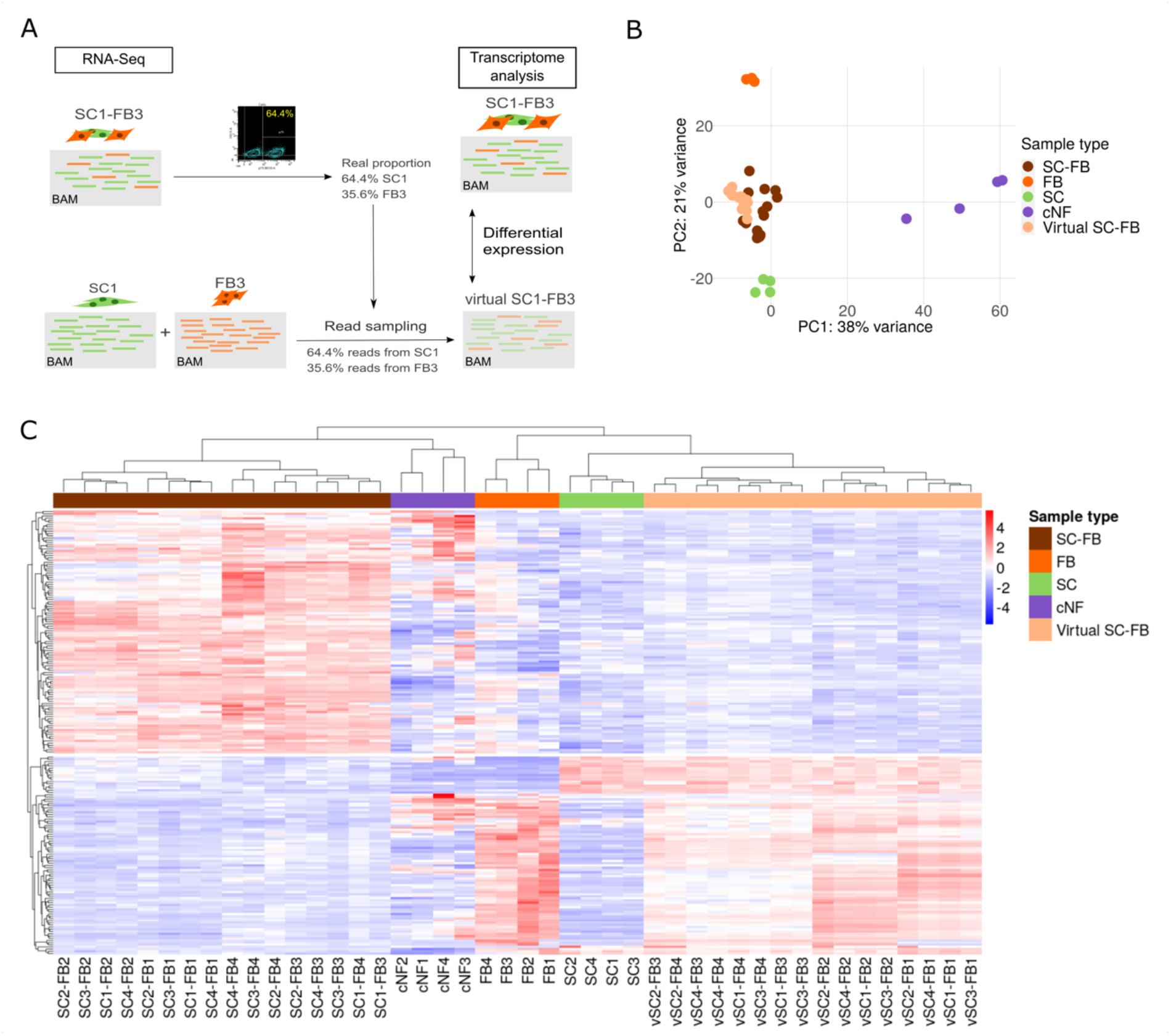
Transcriptome analysis of cNFs, cNF-derived single cultures and SC-FB co-cultures identify expression profiles due to SC-FB interactions. A. Schematic representation of *in-silico* SC-FB co-culture approach to generate virtual co-cultures. Virtual co-cultures were generated by randomly sampling a number of reads from the BAM files of the single cultures, and mixing them from each single culture with the exact proportions of co-cultures. B. Principal component analysis of cNFs, SCs and FBs single cultures, SC-FB co-cultures and virtual SC-FB co-cultures for whole-genome RNA-seq showing the proximity between real and virtual co-cultures. C. Heatmap plot representing an unsupervised cluster analysis of differentially expressed genes (adjusted p-value < 0.05) between real and virtual SC-FB co-cultures. SC: Schwann Cells; FB: Fibroblasts; SC-FB: SC-FB co-cultures, cNF: cutaneous neurofibroma.

**Figure 3.**
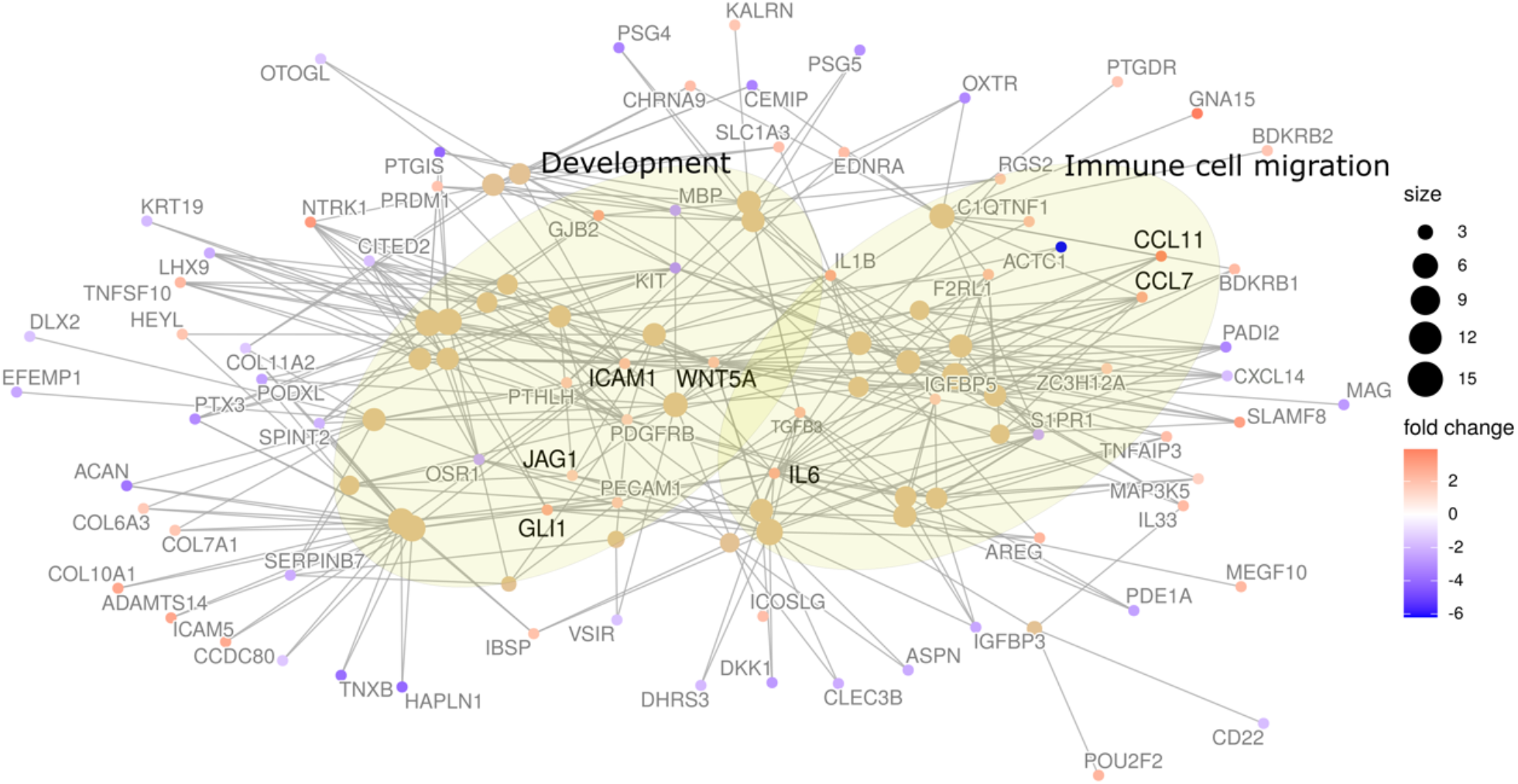
Network analysis of biological processes captured by the differential upregulated genes in real versus virtual SC-FB co-cultures. Network plot showing most differentially upregulated genes and the biological processes (BP) captured by them. Lines define shared genes among the distinct BPs. The size of the sphere representing the BP is proportional to the number of genes contained.

### SC-FB interaction elicits the secretion of multiple cytokines and chemokines involved in immune cell migration

Among the twenty-fifth most significantly enriched biological processes, 9 were related to cytokines and chemokines influencing immune cell regulation and migration (**Figure 4A**). We selected some of the genes representing these biological processes, like CCL5, CCL7, CCL11, ICAM, NCAM, VEGF, and IL6, and performed an expression analysis from RNA-seq data, considering SC and FB single cultures, real and virtual co-cultures and cNFs (**Figure 4B**). All genes showed statistically significant expression differences between real and virtual co-cultures, mainly being upregulated in real co-cultures. To investigate whether transcription differences were also consistent at the protein level, and to investigate their potential paracrine action, we measured the secretion of their protein products in the supernatants of all single SC and FB cultures and all co-cultures using Luminex (see M&M section and **Supplemental File 3**). Again, we generated a companion virtual co-culture for each of the 16 co-cultures in a similar way described in **Figure 2A**, but this time considering the selected protein products being measured in single SC and FB culture supernatants and mixing them according to the real SC-FB proportions in each real co-culture. All secreted products of selected genes were also significantly overexpressed in real vs virtual co-cultures (**Figure 4C**), indicating a correlation from transcriptional up to the secretion of their protein products. Thus, these results indicated that signaling for immune cell recruitment in cNFs is at least partially produced by the SC-FB crosstalk.

**Figure 4.**
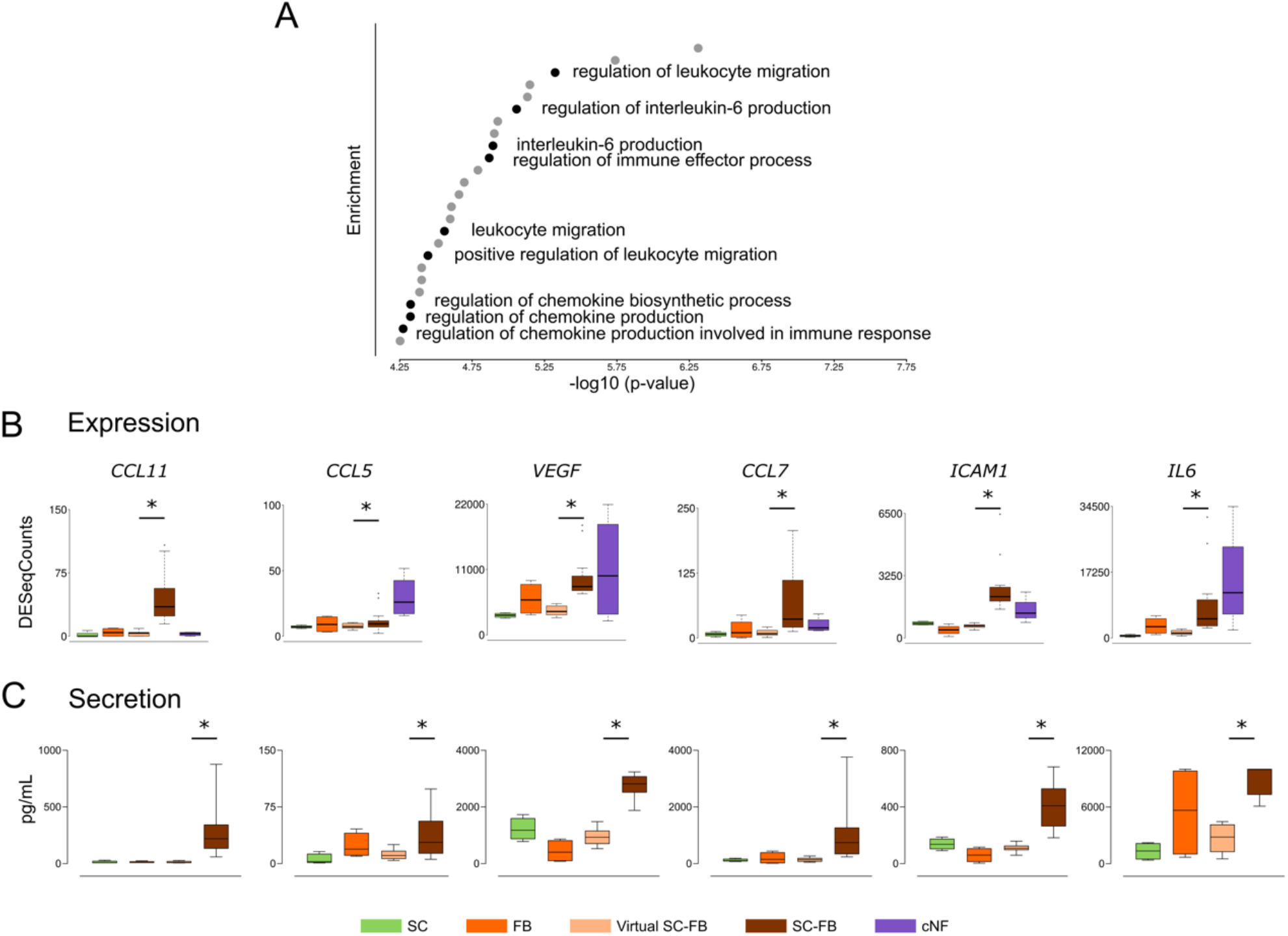
SC-FB interaction elicit the secretion of multiple cytokines and chemokines involved in immune cell migration. A. Enrichment analysis of upregulated genes in real co-culture showing the 25 most significant biological processes (BPs). In black, BPs related to cytokines and chemokines production and immune cell migration are shown. B and C. Correlation between expression data from RNAseq (B) and secretion data from Luminex (C) of several upregulated genes in SC-FB co-culture experiments compared to single cultures and primary tumors. One-tailed paired t-test (p-value ≤ 0.05). SC: Schwann Cells; FB: Fibroblasts; virtual SC-FB: Virtual SC-FB co-cultures; SC-FB: SC-FB co-cultures, cNF: cutaneous neurofibroma.

### Gpr68 activation by Ogerin reduces viability and proliferation of cNF-derived primary SCs in both single cultures and SC-FB co-cultures

We selected a second group of biological processes broadly grouped as involved in developmental processes and developmental signaling pathways. We chose different genes central to these biological processes (**Figure 5; Supplemental Table 1**), like JAG1 (Notch signaling), GLI1 (Sonic Hedgehog signaling); WNT5A (Wnt signaling); TGFb3 (TGFb signaling); AREG (EGF/TGF-alpha signaling); TGFA (TGF-alpha signaling); GPR68 (cAMP signaling). To identify whether any of these pathways had an impact on cNF growth, we selected compounds increasing or decreasing the function of their protein products and performed a preliminary cell viability screening assay using cNF-derived SC cultures (data not shown). From the results obtained, we selected *GPR68, GLI1*, and *WNT5A* as potential targets, to further characterize their effect in SC cultures and SC-FB co-cultures. Expression analysis showed that all three genes were significantly overexpressed in real versus virtual co-cultures (**Figure 5A**). Three different compounds were selected to modulate their gene products: Ogerin, an allosteric activator of the G-protein coupled receptor 68 (Gpr68) which in turn activates adenylyl cyclase and elicits the production of cAMP (**Figure 5Bii**); GANT61, an inhibitor of the transcription factor 1 (GLI1, **Figure 5Biii**); and LGK974, an inhibitor of the secretion of Wnt via Porcupine (**Figure 5Biv**). We used Selumetinib, a MEK inhibitor (MEKi) authorized by the FDA and EMA to treat inoperable pNFs in children (Gross et al., 2020), as a positive control (**Figure 5Bi**). Functional analyses performed and cNF-derived SCs used for each analysis are summarized in **Supplemental Table 2**. We first tested the toxicity of different concentrations of the selected drugs, using NF1 patient-derived skin FBs (**Supplemental Figure 4**), and selected one dose for further experiments (15 μM Ogerin, 5 μM GANT61, and 15 μM LGK974). Next, we characterized the physiological role of these signaling pathways in primary cNF-derived SC and FB single cultures and SC-FB co-cultures. Cells were seeded, and after 24 hours media with the different drugs or vehicle (DMSO) was replaced. We evaluated cell viability through the following 72 hours using a luminometric assay (RealTime-Glo; **Figure 5C**) and cell proliferation at 48 hours using flow cytometry (Click-iT EdU; **Figure 5D-G**) in cultures from three independent cNFs. Selumetinib treatment strongly reduced cell viability in SC cultures and SC-FB co-cultures with a slight effect on cNF-derived FB cultures (**Figure 5C**). A similar viability pattern was observed in Ogerin-treated SC cultures, alone or in co-cultures, with no effect on FBs viability. For GANT61 and LGK974, although exhibiting a certain effect on SC viability, the response on SCs was either not as clear as Ogerin (GANT61) or exhibited inconsistencies among different cNF-derived cells (LGK974). Furthermore, Selumetinib elicited a potent arrest of SCs in single cultures after 48h (**Figure 5D and E**). Using S100B staining in combination with Click-iT EdU we were able to measure independently the proliferation status of SC and FB populations in co-cultures (**Figure 5F and G**). Again, Selumetinib elicited a strong arrest of both SCs and FBs in co-cultures. Ogerin produced a slight arrest on SCs, in both single and co-cultures, that was consistent among cultures derived from the three independent cNFs. For GANT61 and LGK974 responses were either with no effect (LGK974) or inconsistent among biological replicates (GANT61).

**Figure 5.**
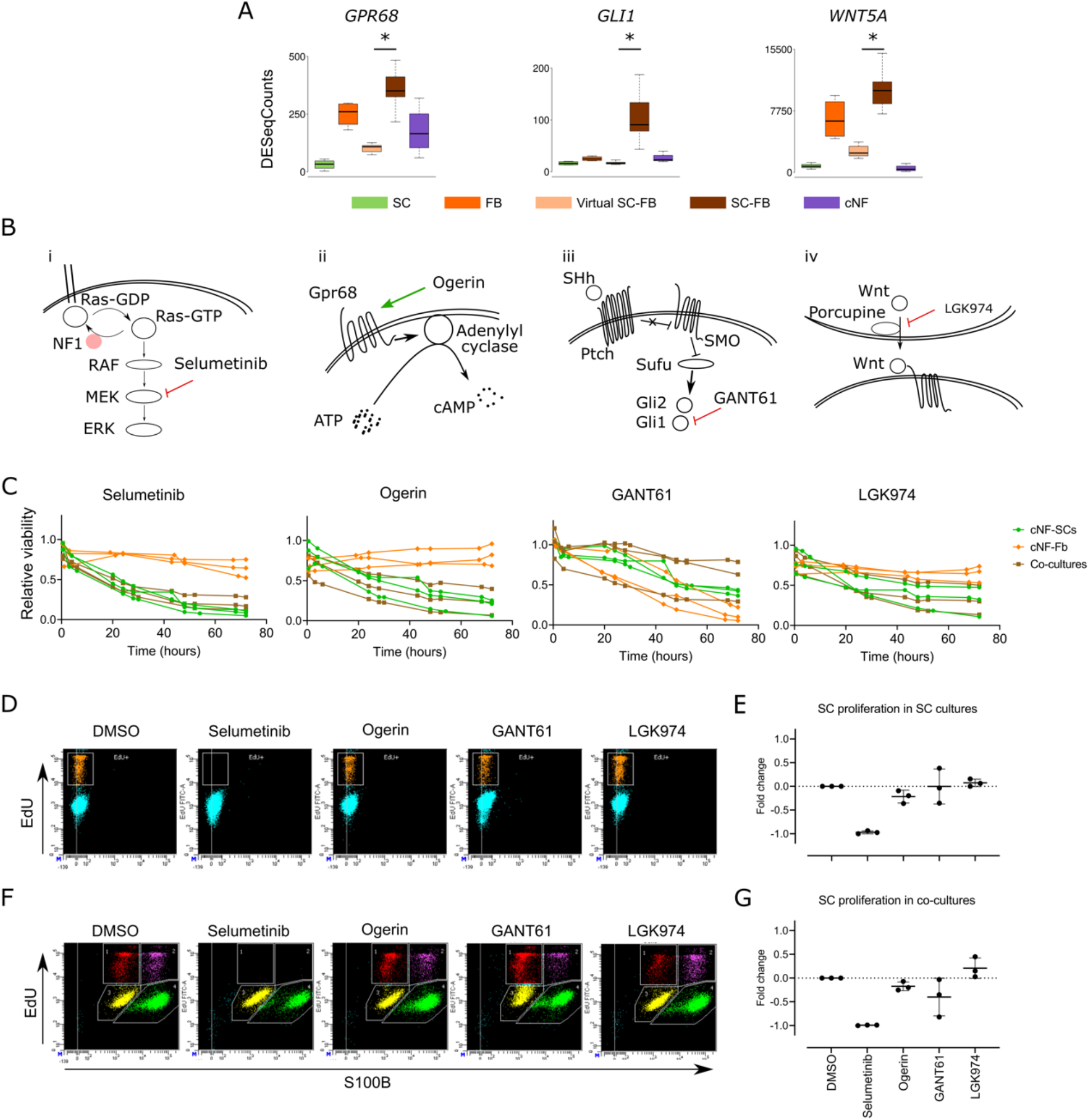
Ogerin decreases SC proliferation and viability in cNF-derived SCs and co-cultures. A. Expression data from RNAseq of *GPR68, GLI1* and *WNT5A* in single cultures, SC-FB co-cultures and cNFs. One-tailed paired t-test (p-value ≤ 0.05 (*)). B. Mechanism of action of selected drugs (i) Selumetinib, (ii) Ogerin, (iii) GANT61 and (iv) LGK974 and related to their transduction signaling pathways. C. Effect of selected drug treatments on cell viability in primary SC (green), FB (orange) and SC-FB co-cultures (brown). Cells were plated and 24h later treated with 40 μM Selumetinib, 15 μM Ogerin, 5 μM GANT61 and 15 μM LGK974. Cell viability was monitored throughout 72h using RealTime-Glo MT Cell Viability Assay. Three different primary samples were used per cell type. Mean is represented. Data are expressed as relative viability to DMSO-treated control cells. D and E. Effect of drug treatments on cell proliferation in primary SC cultures. Cells were plated and 24h later treated with 40 μM Selumetinib, 15 μM Ogerin, 5 μM GANT61 and 15 μM LGK974. After 48 h of treatment cell proliferation was assessed by Click-iT EdU Flow Cytometry Assay. Representative flow cytometry plot of the distinct treatments of one primary SC culture (D). Data are expressed as fold change mean normalized to DMSO-treated control cells ± SEM from three different primary SC cultures (E). F and G. Effect of drug treatments on cell proliferation in SC-FB co-cultures. Cells were plated and 24h later treated with 40 μM Selumetinib, 15 μM Ogerin, 5 μM GANT61 and 15 μM LGK974. After 48 h of treatment cell proliferation was assessed by Click-iT EdU Flow Cytometry Assay in combination with S100B staining to distinguish SC population. Representative flow cytometry plot of the distinct treatments of one SC-FB co-culture (F). Data are expressed as fold-change mean normalized to DMSO-treated control cells ± SEM from three different primary SC cultures (G). SC: Schwann Cells; FB: Fibroblasts; virtual SC-FB: virtual SC-FB co-cultures; SC-FB: SC-FB co-cultures; cNF: cutaneous neurofibroma.

### Selumetinib and Ogerin co-treatment induces loss of viability, differentiation, and death of *NF1* (-/-) SCs

MEKis treatment results on plexiform neurofibromas (Dombi et al., 2016; Gross et al., 2020; Jessen et al., 2013) opened the possibility of using these drugs likewise for cNFs, either alone or potentially combined with other agents. Given the role of Ras and cAMP pathways in the balance of SC proliferation/differentiation, we hypothesized that unbalancing both pathways by the Selumetinib-Ogerin co-treatment could serve for preventing *NF1*(-/-) SC proliferation in cNFs. To test this possibility, we first confirmed that Ogerin treatment was elevating cAMP levels in cNF SCs (**Supplemental Figure 5**). Then, we examined the toxicity effect of Selumetinib-Ogerin co-treatment on NF1 skin-derived FBs, monitoring viability by RealTime-Glo for 72h (**Supplemental Figure 6**). This co-treatment elicited no toxicity on skin FBs, either using low or high Selumetinib concentrations. Next, we tested the co-treatment effect on *NF1*(-/-) SC viability in three independent cNF SC cultures, employing the same assay (**Figure 6A**). Selumetinib-Ogerin co-treatment, using either low or high concentrations of Selumetinib, enhanced the loss of viability of SCs compared to single agents alone. To broader the analysis of SC physiology, we carefully characterized the impact of treatments on SC morphology. We stained SC cultures with S100B. While Ogerin alone elicited no apparent morphological change in SCs, Selumetinib, especially at high concentrations, enhanced the rectilinear and spindle shape morphology of SCs, decreasing the cytoplasmic volume (**Figure 6B**). In a completely opposite direction, Selumetinib-Ogerin co-treatment made SCs spread, flatten, and expanded the cell membrane (**Figure 6B**). To rule out whether the morphological change of co-treated SCs was reflecting SC differentiation and myelinization, we immuno-stained these cells using the myelin protein zero (MPZ) antibody. While cells treated with single agents alone did not stain for MPZ protein compared to controls, co-treated cells were highly MPZ positive, indicating a pro-myelinization differentiating SC phenotype. To explore if this physiological change was specific to the GPR68 activation or was produced by the activation of the cAMP pathway, we substituted Ogerin with the cAMP analog 8-CPT cAMP. This agent alone elicited no change in MPZ expression in SCs, but when combined with Selumetinib, again cNF SCs adopted a spread and flatten phenotype, generating also neurites, and expressed high quantities of MPZ (**Figure 6C**). This result indicated that the SC differentiation phenotype was dependent on cAMP elevation, through Gpr68 activation or independently of it. We observed that in addition to differentiating SCs, co-treated SC cultures contained a significant number of dead cells and cell debris in the cultures. Thus, we measured apoptosis using flow cytometry analysis (**Figure 6D and Supplemental Figure 7**). Ogerin alone almost had no effect on SC apoptosis, but Ogerin-Selumetinib co-treatment enhanced the degree of apoptosis compared to Selumetinib as a single agent, both at low and high concentrations of the MEKi. Altogether, Ogerin-Selumetinib co-treatment induced loss of viability, SC myelinization, and death of cNF-derived primary SCs.

**Figure 6.**
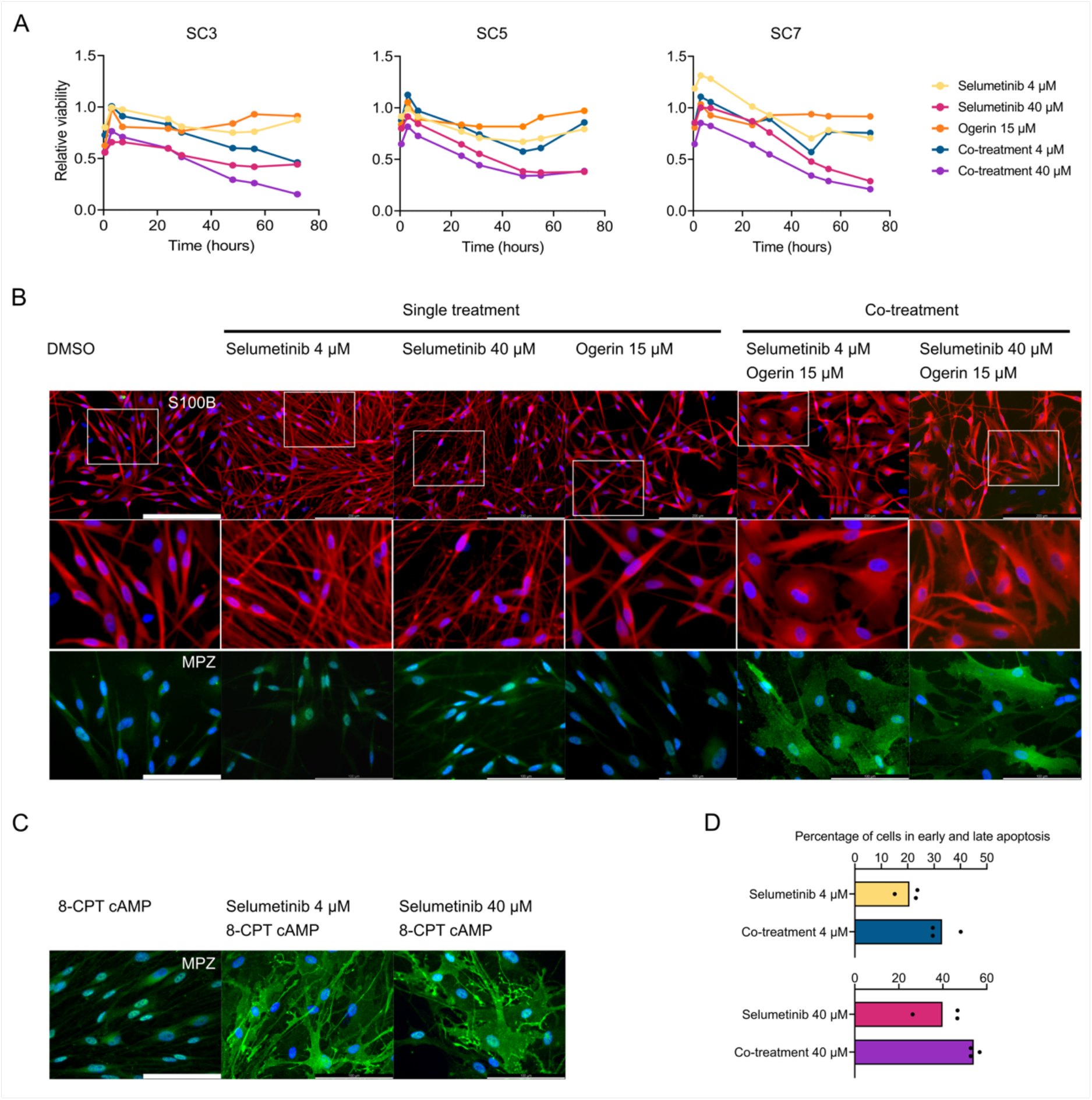
Selumetinib and Ogerin co-treatments lead to SC differentiation and increased cell death in cNF-derived SCs. A. Effect of Selumetinib and Ogerin treatments and co-treatments on cell viability in primary SC cultures. Cells were plated and 24h later treated with single drugs (4 μM Selumetinib, 40 μM Selumetinib, 15 μM Ogerin) or as co-treatment of drugs (4 μM Selumetinib and 15 μM Ogerin; 40 μM Selumetinib and 15 μM Ogerin). Cell viability was monitored throughout 72h using RealTime-Glo MT Cell Viability Assay. Three different primary SC cultures were used (SC3, SC5, SC7). Mean is represented. Data are expressed as relative viability to DMSO-treated control cells. B. Immunocytochemical analysis for S100B and MPZ of treated SC cultures. DAPI was used to stain cell nuclei. Scale bars, 200 μm (S100B) except in enlarged view; 100 μm (MPZ) C. Immunocytochemical analysis for MPZ of SC cultures treated with the cAMP analogue 8-CPT (250 μM) in combination with Selumetinib (4 μM and 40 μM). DAPI was used to stain cell nuclei. Scale bars, 100 μm. D. Effect of Selumetinib and Ogerin treatments and co-treatments on cell death in primary SC cultures. Cells were plated and 24h later treated with single drugs (4 μM Selumetinib, 40 μM Selumetinib, 15 μM Ogerin) or as co-treatment of drugs (4 μM Selumetinib and 15 μM Ogerin; 40 μM Selumetinib and 15 μM Ogerin). Cell death was monitored using the Flow cytometry Annexin V Apoptosis Detection Kit. Data are expressed as the percentage of cells in early and late apoptosis per condition in three different primary SC cultures. One-tailed unpaired t-test (p-value ≤ 0.05 (*)).

### Co-treatment of Selumetinib and Ogerin induces sphere disaggregation and cell death in an iPSC-derived 3D neurofibromasphere model

We decided to validate Selumetinib-Ogerin co-treatment results in an independent neurofibroma model that we recently developed (Mazuelas et al. 2022). This model consists of the generation of neurofibromaspheres, composed of iPSC-derived *NF1*(-/-) SCs (∼70%) mixed with neurofibroma-derived FBs (∼30%). Upon engraftment in the sciatic nerve of nude mice, these neurofibromaspheres generate genuine neurofibroma-like tumors, with histology recapitulating most features of human neurofibromas (Mazuelas et al., 2022). For this experiment, we used neurofibromaspheres containing cNF-derived FBs together with iPSC-derived *NF1*(-/-) differentiating SCs (**Figure 7A**). Single treatment and co-treatments were evaluated after 72h. At this point, spheres under co-treatment conditions lost compactness and exhibited an enhanced disaggregated appearance compared to single treatments and controls (**Figure 7B**). We quantified disaggregation and generated a disaggregation index (see M&M section; **Figure 7C**). Selumetinib-Ogerin co-treatments exhibited a significantly enhanced disaggregation index compared to single agents and controls. Furthermore, by using a life and death staining assay, we observed a significantly increased normalized death signal in Selumetinib-Ogerin co-treatments, compared to single treatments alone, especially at high concentrations of Selumetinib (**Figure 7D**). Significant differences were evidenced after quantification (**Figure 7E**). Thus, Selumetinib-Ogerin co-treatment response on neurofibromaspheres was consistent with results obtained using cNF-derived primary SC cultures and SC-FB co-cultures, pointing to a potential therapeutic strategy for cNFs.

**Figure 7.**
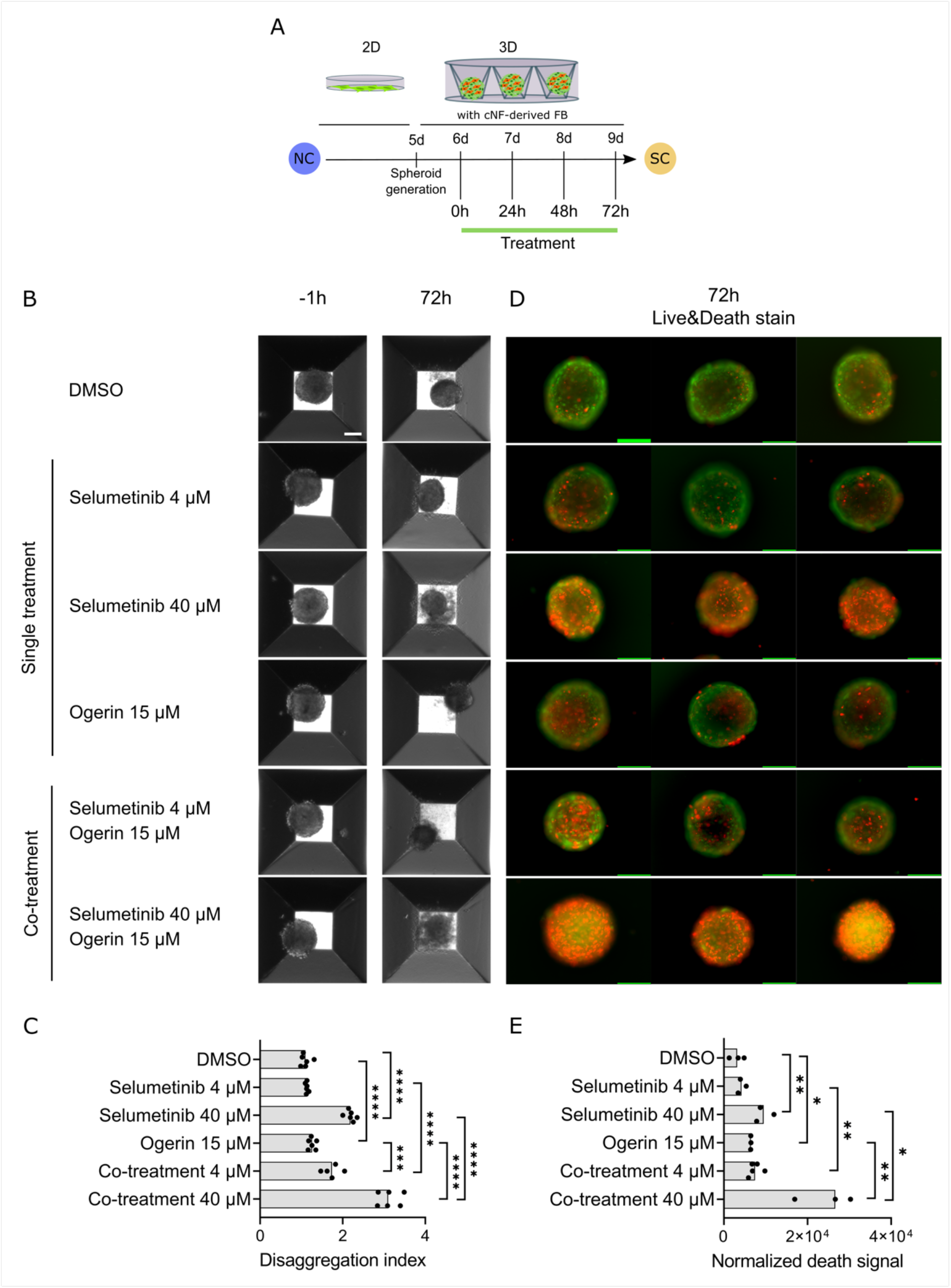
Selumetinib and Ogerin co-treatments increased cell death in an iPSC-based SC-FB cNF model in 3D. A. Schematic representation of drug treatment in iPSC-based SC-FB cNF model in 3D (neurofibromaspheres). *NF1*(-/-) iPSC-derived NC cells were plated in SC differentiation media for 5 days. At this point, cells were mixed with cNF-derived FB and plated in microcavity Aggrewell plates to generate neurofibromaspheres. 24h after seeding, neurofibromaspheres were treated with single treatments (4 μM Selumetinib, 40 μM Selumetinib, 15 μM Ogerin) or co-treatments (4 μM Selumetinib and 15 μM Ogerin; 40 μM Selumetinib and 15 μM Ogerin) for 72h. B. Representative phase-contrast images showing differences in neurofibromasphere morphology at 72h between vehicle DMSO-treated spheroids, single treatment and co-treatments. Scale bar: 100μm. C. Disaggregation index (DI) of neurofibromaspheres showing a higher effect in co-treatments than in single treatments. DI is calculated by measuring disaggregated spheroid area normalized by spheroid area at 72h of drug treatment. Data is shown as the median of n ≥ 5 spheroids. One-tailed unpaired t-test (p-value ≤: 0.05 (*), 0.01 (**),0.001 (***), and 0.0001 (****). D. Representative images of Acridine Orange (Live, green) and Propidium iodide (Death, red) stained neurofibromaspheres after 72h of drug treatment. Scale bars,100 μm. E. Normalized death signal (NDS) of neurofibromaspheres showing a higher effect in co-treatments than in single treatments. NDS is calculated as Propidium Iodide intensity normalized by the area generated in Acridine Orange channel at 72h of drug treatment. Data is shown as the median of n ≥ 3 spheroids. One-tailed unpaired t-test (p-value ≤: 0.05 (*), 0.01 (**),0.001 (***), and 0.0001 (****). FB: fibroblast; iPSC: induced pluripotent stem cells; NC: neural crest; SC: Schwann cell.

## Discussion

We set up a robust experimental framework that was carefully executed to capture the crosstalk between SCs and FBs at a transcriptome level. The transcriptome of 16 SC-FB co-cultures was analyzed and compared to that of their corresponding single cultures and original cNFs, to minimize the potential biological biases of particular cNFs or cell types. Indeed, the unsupervised cluster analysis of the transcriptome of SC-FB co-cultures evidenced the dominant influence of FBs over SCs in the whole expression signature (**Figure 2C**). The identified SC-FB crosstalk signature representing overexpressed genes was also mostly preserved in an iPSC-derived mixed SC-FB neurofibromasphere model (Mazuelas et al., 2022) and not in SC-only spheres, strongly supporting the SC-FB interaction nature of the obtained expression signature. Enrichment analysis of this signature highlighted the presence of biological processes components related to immune response, immune cell migration, and chemotaxis; and also signaling pathways important in developmental processes.

The most studied component of the cNF microenvironment has been the immune infiltrate (revised in (Fletcher et al., 2020)). Different molecules have been implicated in the recruitment of different cell types (mast cells, macrophages, T-cells, dendritic cells) to neurofibromas: CCL2/CCL12/CCR2 (Fletcher et al., 2019a); CSF1 (Prada et al., 2013); CXCL10/CXCR3 (Fletcher et al., 2019b); IL-1b (Choi et al., 2017); IL-6 (Wu et al., 2017); KITLG (SCF) (Liao et al., 2018; Yang et al., 2003). In the present work, we identified some of these genes to be transcriptionally upregulated due to the SC-FB interaction (CCL2, CXCL10, IL1B, IL6) and not others (CSF1), or to a lesser extent (CCL5, KITLG). CSF1 and KITLG were highly expressed in FBs. At the protein level, secreted in supernatants of SC-FB co-cultures, we confirmed the upregulated secretion of IL6, CCL5, and other factors such as CCL11, CCL7, ICAM, and angiogenic factors such as VEGF (**Supplemental File 3**). In this regard, our works shows that at least part of the secreted proteins that promote immune cell recruitment into cNFs is specifically due to the interaction between SCs and FBs.

Components of signaling pathways involved in developmental processes were also enriched, like JAG1 (Notch), GLI1 (Sonic Hedgehog); WNT5A (Wnt); TGFb3 (TGFb); AREG (EGF/TGF-alpha); TGFA (TGF-alpha); GPR68 (cAMP). We performed preliminary assays with all of them and viability and proliferation assays in cNF-SCs after the alteration by drug treatment of a few selected components. We identified the receptor GPR68 as an interesting target for cNF biology and/or treatment. We used Ogerin, an allosteric modulator of GPR68 that potentiates proton activity and activates adenylyl cyclase, to activate GPR68. Upon Ogerin treatment, cNF-derived SCs increased the levels of cAMP, decreased SC viability, and slightly decreased SC proliferation in single cultures and SC-FB co-cultures. In contrast, Ogerin had no effect on cNF-derived FBs, or on FBs derived from the skin of an NF1 individual. Ogerin has already been used *in vitro* and *in vivo* (Huang et al., 2015) but not much information has been generated regarding its potential off-target effects, toxicity, kinetics, and dynamics, using *in vivo* models. Other GPR68 activators have been developed, like a much more potent allosteric activator (∼30 fold) termed MS48107 (or PAM71) based on Ogerin structure (Yu et al., 2019). In addition, three positive allosteric peptides have been identified (Osteocrin^33-55^, CART(42-89)^9-28^, and corticotropin^17-40^) with up to 2-fold improved allosteric activity compared to Ogerin (Foster et al., 2019), providing alternatives to the use of Ogerin for the activation of GPR68. We observed that Ogerin increases the intracellular levels of cAMP in cNF-derived SC cultures (**Supplemental Figure 5**).

The discovery of MEKis to treat plexiform neurofibromas opened the possibility to use these drugs likewise for cNFs (Dombi et al., 2016; Gross et al., 2020; Jessen et al., 2013). However, despite effectiveness, results with pNFs and first results using the *Nf1*-KO model (*Prss56*^Cre^, *Nf1*^fl/fl^) (Coulpier et al., 2022) indicated that MEKi alone might not be a complete solution for mature cNFs, and the combination of MEKi with other agents emerged as a logical next step for cNF treatment. Neurofibromin plays a role in SCs and neurofibromas by regulating both the Ras/MAPK pathway (Kim et al., 1995; Sherman et al., 2000) and the cAMP pathway (Kim et al., 2001; Patritti-Cram et al., 2022; Serra et al., 2000). Both pathways need to be active for a prolific SC proliferation (Rahmatullah et al., 1998). In fact, we collected evidence that the regulation of both pathways by neurofibromin could be coordinated (Biayna et al., 2021). However, it is known that in SCs the level of cAMP signaling is crucial for switching from a proliferative state, to a myelin-forming differentiation state (Arthur-Farraj et al., 2011). Thus, with all this information, and the results obtained from exposing cNF-derived SCs to Ogerin, we reasoned that the unbalancing of both pathways by the simultaneous combination of decreasing the RAS pathway and increasing the cAMP pathway could be a good strategy for preventing *NF1*(-/-) SC proliferation and potentially a therapeutic strategy for cNFs. Thus, Selumetinib-Ogerin co-treatment enhanced the loss of viability of SCs compared to single agents alone. In addition, it elicited a pro-myelinization differentiating SC phenotype, also reproduced by replacing Ogerin with a cAMP analog, indicating a cAMP-dependent effect. Finally, co-treatment enhanced apoptosis of SCs compared to Selumetinib as single agent. These results not only support this co-treatment as a potential strategy for cNFs but represent the first *in vitro* data of the effect of Selumetinib alone or in combination using human cNF-derived primary SCs and FBs. The corroboration of these results in the orthogonal iPSC-derived 3D neurofibromasphere model (Mazuelas et al., 2022) provides additional evidence, in a relevant physiological model, of the potential of this co-treatment strategy.

Ogerin has already been used *in vivo* but with a brief temporal window (Huang et al., 2015). There is not much information on the toxicity and potential use of Ogerin *in vivo* in treatments involving a prolonged time, such as those that would be needed for treating cNFs. However, as explained before, other GPR68 activators have been developed, with a higher allosteric power for activating GPR68 (Yu et al., 2019) or with a peptide nature (Foster et al., 2019). It would be worth to explore also their possibilities both *in vitro* and *in vivo* in the context of SCs and cNFs. Furthermore, the striking induction of SC differentiation by using the cAMP analog 8-CPT cAMP in combination with Selumetinib supported the cAMP signaling involvement in the results obtained and opened the possibility of using other cAMP activators, independent of GPR68. The cAMP signaling pathway is complex (Sassone-Corsi, 2012), containing many activating and inhibiting components, with different isoforms for each component, like adenylyl cyclases, G-protein coupled receptors; G-alpha -beta and -gamma subunits; phosphodiesterases; etc. The functioning of the pathway in a given cell type is greatly controlled by the abundance of specific isoforms and their regulation (Emery et al., 2015; Hanoune et al., 2001), that at the same time, could be differentially perturbed by specific compounds (Hayashi et al., 1998). In this work, and others (e.g. (Gosline et al., 2017)) extensive-expression data on cNFs and derived cells have been produced to explore the exact isoform spectrum of cAMP-components in *NF1*(-/-) SCs and other cNF cell types. This analysis should allow the identification of other specific cAMP-pathway activators in addition to Ogerin and related compounds, and also provide a layer of specificity to the target cells to decrease potential toxicity effects.

In summary, we identified GPR68 as one of the upregulated genes within a transcriptional signature specifically produced by the interaction of cNF-derived SCs and FBs in co-cultures. The GPR68 allosteric activator Ogerin affected the viability of cNF-SCs in single cultures and SC-FB co-cultures, opening the possibility of exploring a co-treatment with Selumetinib. Thus, we tested the hypothesis that the concomitant inactivation of the Ras/MAPK pathway and the activation of the cAMP pathway may prevent *NF1*(-/-) SC proliferation. Indeed, the Selumetininb-Ogerin co-treatment reduced viability, induced differentiation, and death of cNF-SCs, while not perturbing skin fibroblasts from NF1 individuals. These results were corroborated in an iPSC-derived 3D neurofibromasphere model. The promising results obtained in this work open the possibility of using the unbalancing of the Ras/MAPK and cAMP/PKA pathways by a MEKi and a cAMP elevator co-treatment, as a therapeutic strategy for cNFs.

## Materials and Methods

### Patients, Cutaneous Neurofibromas (cNFs), and tumor processing

NF1 patients diagnosed according to standard diagnostic criteria (Legius et al., 2021) kindly provided tumor samples after giving written informed consent. Immediately after excision, tumor samples were placed in DMEM medium (Gibco) containing 10% FBS (Gibco) + 1x GlutaMax (Gibco) + 1x Normocin antibiotic cocktail (InvivoGene) and shipped at room temperature to our laboratory. Tumors were processed as follows: surrounding fat tissue and skin were removed, and tumors were cut into 1-mm pieces and cryopreserved in 10% DMSO (Sigma) + 90% FBS (Gibco) until used.

### cNF-derived Schwann cells and endoneurial Fibroblasts cultures

CNF-derived Schwann cells (SCs) and endoneurial fibroblasts (FBs) were isolated as described previously (Serra et al., 2000). Briefly, cryopreserved cNFs were thawed, cut into smaller pieces using a scalpel, and digested with 160 U/mL Collagenase Type 1 and 0.8 U/mL Dispase (Worthington, Lakewood, NJ) for 16 hours at 37ºC, 5% CO_2_.

To establish SC cultures, dissociated cells were seeded onto 0.1 mg/mL Poly-L-lysine (Sigma) and 4 μg/mL Laminin (Gibco)-coated dishes in Schwann Cell Media (SCM) which is DMEM (Gibco) with 10% FBS (Gibco), 100 U/mL Penicillin/100 mg/mL Streptomycin (Gibco), 0.5 mM 3-iso-butyl-1-methilxantine (IBMX, Sigma), 2.5 μg/mL Insulin (Sigma), 10 nM Heregulin-b1 (PeproTech), and 0.5 μM Forskolin (Sigma).

Once the SC culture was established, cells were passaged when they reached confluency with 0.05% Trypsin-Ethylenediaminetetraacetic acid (EDTA) (Gibco) and plated in SCM. 24 hours later, the culture medium was replaced by SCM without Forskolin, and 24 hours later media was changed and replaced for SCM for an additional 2–3 days. This process was repeated in cycles. Cells were maintained at 37ºC under a 10% CO_2_ atmosphere. **Supplemental Table 2** summarizes all cNF-derived SCs used for each of the distinct functional analyses.

To establish FB cultures, tumor-dissociated cells were plated in DMEM supplemented with 10% FBS media and 1x GlutaMAX (Gibco) and 100 U/mL/Penicillin/100 mg/mL Streptomycin (Gibco). Cells were passaged when necessary with 0.25% Trypsin-EDTA and maintained at 37ºC under a 5% CO_2_ atmosphere.

### Schwann cell and endoneurial fibroblast co-culture experiment

70% SCs and 30% FB co-cultures were grown under SC culture conditions with some modifications: cells were seeded onto 0.1 mg/mL Poly-L-lysine (Sigma) and 4 μg/mL Laminin (Gibco)-coated dishes in SCM without IBMX (Sigma) to favor FB growth. A total of 5 × 10^4^ cells per cm^2^ were seeded, and 24 hours later, the media was changed to SCM without IBMX nor Forskolin. Co-cultures were maintained at 37ºC under a 10% CO_2_ atmosphere for a total of 72 hours. At this point, supernatants were collected for further analysis, and cells were trypsinized using 0.25% Trypsin-EDTA. A fraction of cells was analyzed for p75 expression by flow cytometry, and another fraction was pelleted and frozen for RNA extraction.

### Neurofibromasphere generation

Neurofibromaspheres were prepared following the protocol described in Mazuelas et al. (2022) using cNF-derived FBs.

### DNA extraction

Genomic DNA from tumors was extracted using the Gentra Puregene Kit (Qiagen, Chatsworth, CA) following the manufacturer’s instructions, after homogenization using Tissue Lyser (Qiagen). Genomic DNA from primary cells was extracted using Promega Maxwell 16 system following manufacturer’s instructions.

### RNA extraction

Tumors were thawed in DMEM supplemented with 10% FBS, and homogenized using TissueRuptor II (Qiagen), and total RNA was extracted using TriPure Isolation Reagent (Roche) following the manufacturer’s instructions. Total RNA from primary cells was extracted using the 16 LEV simplyRNA Purification Kit (Promega) following manufacturer’s instructions in the Maxwell 16 Instrument (Promega). RNA was quantified with Nanodrop 1000 spectrophotometer (Thermo Scientific). Quality was assessed with Bioanalyzer 2200 TapeStation (Agilent).

### NF1 genetic analysis

*NF1* germline and somatic mutations were detected by *NF1* gDNA sequencing using the I2HCP NGS custom panel (Castellanos et al., 2017). Germline mutations were confirmed by DNA Sanger sequencing from cultured cNF-derived FB cells. Loss of heterozygosity of *NF1* locus was detected by Microsatellite Multiplex PCR Analysis (MMPA) of chromosome 17 (Garcia-Linares et al., 2012). The reference sequence used was NG_009018_1; NM_000267_3; NP_000258.1. For intragenic deletions we used NM_001042492.2.

### Exome sequencing

The exome was captured using Agilent SureSelect Human All Exon V5 kit (Agilent, Santa Clara, CA, US) and sequenced in a HiSeq instrument (Illumina, San Diego, CA, US) at Centro Nacional de Analisis Genomicos (CNAG, Barcelona, Spain). Sequencing reads were then mapped with bwa mem (Li, 2013) against GRCh38. We called variants with strelka2 (Kim et al., 2018) and annotated them with annovar (Wang et al., 2010).

### RNAseq analysis

RNA-seq libraries were prepared in the IGTP Genomics Core Facility using the TruSeq stranded mRNA Illumina, quantified with the KAPA library quantitation kit for Illumina GA and the Agilent Bioanalyzer, and sequenced at CNAG in a HiSeq platform pooling 3 samples per lane (paired-end, 2&100).

To evaluate the impact of the heterotypic interactions on the co-culture transcriptomic profiles we created a matched set of virtual co-cultures by *in-*silico mixing sequencing reads from the pure SC and FB. To build them, we randomly sampled a total of 38.5 million reads from the fastq files of the SC and FB in each co-culture in the exact same proportion of SC and FB determined by flow cytometry analysis. These reads were then stored in a new fastq file representing the expected transcriptional profile of each co-culture if no heterotypic interactions were present. The differential expression analysis between each real co-culture and its virtual counterpart revealed the transcriptional changes due to heterotypic interactions. RNA-seq data from real and *in-*silico mixed samples was aligned with Salmon v1.8.0 (Patro et al., 2017) against the reference genome and transcriptome (refMrna and hg38 genome from UCSC)(Patro et al., 2017)(Patro et al., 2017)(Patro et al., 2017)(Patro et al., 2017)(Patro et al., 2017)(Patro et al., 2017). We imported transcript-level estimates into R and summarized them to gene-level using tximport (Soneson et al., 2016). We then filtered out genes with less than 5 counts in more than two samples and used DESeq2 (Love et al., 2014) to perform differential expression analysis. We finally used clusterProfiler (Yu et al., 2012) to determine Gene Ontology (GO) and KEGG pathways enrichment from differentially expressed genes (p-adjusted value < 0.05). Hetmaps were created using the heatmap R package.

### Flow cytometry of p75 in single cNF-derived cultures and cocultures

For flow cytometry analysis of p75 in primary single SC and FB cultures and SC-FB cocultures, cells were detached with 0.25% Trypsin-EDTA, washed with 1% BSA (Sigma) in PBS, incubated for 30 minutes on ice with unconjugated primary antibody p75 (1:1000, Ab3125, Abcam), washed with 1% BSA in PBS and incubated with Alexa Fluor 488-conjugated secondary antibodies 1:1000 for 30 minutes on ice. Cells were analyzed by flow cytometry using BD LSR Fortessa SORP and BD FACSDiva 6.2 software.

### Immunofluorescence

Cells were fixed in 4% paraformaldehyde (Chem Cruz) in PBS for 15 minutes at room temperature, permeabilized with 0.1%Triton-X100 in PBS for 10 minutes, blocked in 10% FBS in PBS for 15 minutes, and incubated with primary antibodies, p75 (1:100, Ab3125, Abcam), S100B (1:1000, Z0311, Dako) and vimentin (1:200, MA5-11883, ThermoFisher) overnight at 4ºC. Secondary antibodies were Alexa Fluor 488- and Alexa Fluor 568-(Thermo Fisher Scientific). Nuclei were stained with DAPI (Stem Cell Technologies, 1:1000).

### Measurement of cytokines and chemokines in co-culture supernatants

Supernatants were collected 72 hours post-seeding the cells, transferred into clean polypropylene microcentrifuge tubes, and centrifuged at 14,000 rpm for 10 minutes at 4ºC to remove any cellular debris. The clarified mediums were aliquoted into clean polypropylene microcentrifuge tubes and stored at -80ºC until used.

On the day of the analysis, frozen samples were thawed on ice and centrifugated at 14,000 rpm for 10 minutes at 4ºC. Selected cytokines and chemokines including BDNF, sICAM-1, and NCAM (Cat. No HNDG3MAG-36K), S100B and GDNF (Cat. No HCYTOMAG-60K), and EGF, FGF2, Eotaxin, TGFα, G-CSF, fractalkine, MCP-3, PDGF-AA, IL-1α, IL-1β, IL-6, IL-8, CXCL10, MCP-1, MIP-1α, RANTES, and VEGF (Cat. No HCYTOMAG-60K) were quantified using Milliplex MAP Luminex microbead assays according to the manufacturer’s instructions (Merck Millipore, Darmstadt, DE). Samples were analyzed without dilution in duplicate, and plates were analyzed on a Luminex 200 with xPONENT software (Luminex Corp., Texas, USA).

### Drug treatment

Drugs used in this study were purchased from commercial sources: Selumetinib (Tocris Cat num. 6815); Ogerin (Tocris Cat num. 5722); GANT61 (Tocris Cat num. 3191); LGK-974 (Selleckchem Cat num. S7143) and prepared as indicated from the manufacturer. For drug treatments, cells were seeded either onto 0.1 mg/mL Poly-L-lysine (Sigma) and 4 μg/mL Laminin (Gibco)-coated plates (SCs and co-cultures) or in non-coated plates (FBs) and 24 h later drugs were added at the indicated concentration. A vehicle-treated control (DMSO) was applied to each cell type at the same dilution used to deliver the drug.

### Cell Viability Analysis

Cell viability in primary SCs, FBs, and co-cultures was assessed using the RealTime-Glo MT Cell Viability Assay (Promega). Briefly, cells were seeded either onto 0.1 mg/mL Poly-L-lysine (Sigma) and 4 μg/mL Laminin (Gibco)-coated opaque 96-well plates in SCM (SCs and co-cultures) or non-coated opaque 96-well plates in DMEM 10%FBS (FBs) at a density of 500 cells/well. 24 h later drugs were added in SCM w/o IBMX nor Forskolin (SCs and co-cultures) or DMEM 10%FBS (FBs) together with the RealTime -Glo reagent following manufacturer’s instructions. Luminescence was monitored for 72 h on a Varioskan Flash plate reader (Thermo). Cells from three independent patients were used (see **Supplemental Table 2**).

### Cell Proliferation Analysis

Cell proliferation was assessed using the Click-iT EdU Flow Cytometry Assay Kit (Thermo Fisher). One hundred thousand primary SCs from 3 independent patients (see **Supplemental Table 2**) were plated onto 0.1 mg/mL Poly-L-lysine (Sigma) and 4 μg/mL Laminin (Gibco)-coated 12-well plates in SCM without IBMX. 24 h later drugs were added in SCM without IBMX nor Forskolin. 48 h after drug treatment cells were treated with 5 μM EdU for 16 hours, fixed, permeabilized, and click labeled with Alexa Fluor 488 azide according to the manufacturer protocol. Cells were also stained with propidium iodide to detect DNA content. Data was collected and analyzed using an FACSCanto II (BD Biociencias) and BD FACSDiva 6.2 software.

### Apoptosis Analysis

One hundred thousand primary SCs from 3 independent patients (**see Supplemental Table 2**) were plated onto 0.1 mg/mL Poly-L-lysine (Sigma) and 4 μg/mL Laminin (Gibco)-coated 12-well plates. 24 h later drugs were added in SCM w/o IBMX nor Forskolin. 48 h after drug treatment, cells were harvested using trypsin-EDTA 0,05%, washed in PBS, and incubated with Annexin-V-FITC antibody using the Annexin V FITC Apoptosis detection kit (Invitrogen) following the manufacturer’s instructions. Data was collected and analyzed using a FACSCanto II (BD Biociencias) and BD FACSDiva 6.2 software.

### Quantification of cAMP levels

Intracellular cAMP levels were quantified using the cAMP-Glo Max Assay (Promega) following manufacture’s instructions. Five thousand cNF-derived primary SCs from 3 independent patients (see **Supplemental Table 2**) were plated onto 0.1 mg/mL Poly-L-lysine (Sigma) and 4 μg/mL Laminin (Gibco)-coated opaque 96-well plates in SCM without IBMX with Forskolin and maintained at 37ºC under a 10% CO2 atmosphere for 24 hours. At this point, cells were treated with an induction buffer containing Ogerin, Forskolin (as a positive control), or neither of them for 30 minutes at room temperature, and Luminescence was monitored on a Varioskan Flash plate reader (Thermo). The induction buffer was composed of PBS w/o calcium nor magnesium, 500 μM IBMX, 100 μM Ro 20-1724 (Sigma, B8279), and 25 mM MgCl_2_.

### Quantification and statistical analysis

Data were analyzed and graphically represented using Microsoft Office Excel spreadsheet and GraphPad Prism (version 9.4.1). Quantitative data are shown as the mean ± standard error (SEM) of three independent experiments.

Bioinformatic analysis of RNA-seq is thoroughly described in the “*RNA-Seq and analysis*” in the method details section, including the exact software and statistical methods used. We applied the default multiple testing correction recommended by the different software packages when applicable. The significance level was established at p < 0.05 except if otherwise stated.

The statistical analysis of Apoptosis data was conducted using a one-tailed unpaired t-test (p-value ≤ 0.05).

For neurofibromasphere analysis, disaggregation index (DI) and Normalized death signal (NDS) were calculated using Image J software. DI was calculated by measuring disaggregated spheroid area normalized by spheroid area at 72h of treatment and co-treatments. Data are shown as the median of n ≥ 5 spheroids. One-tailed unpaired t-test (p-value ≤ 0.05). NDS was calculated as Propidium Iodide intensity normalized by the area generated in the Acridine Orange channel at 72h of treatment and co-treatments. Data are shown as the median of n ≥ 3 spheroids. One-tailed unpaired t-test (p-value ≤ 0.05).

The meaning of the value of n, and/or dispersion and precision measure (SEM) can be found in the Figure legends and Results section.

### Study approval

This study was revised and approved by the IGTP Institutional Review Board. Written informed consent was received from all NF1 patients that provided tumor samples prior to participation in this study.

## Author contribution

Designing research studies (HM, MC, BG, ES); conducting experiments (HM, IU-A, MC); bioinformatic design and analysis (MM-L, BG); acquiring and analyzing genetic data (HM, AN, IR, IB, EC, CL); providing reagents (CL); generation of figures (HM, MM-L, MC, BG, ES); writing the manuscript (HM, MC, ES); Review & Editing (HM, MM-L, IU-A, …, MC, BG, ES).

## Acknowledgements

This work has mainly been supported by an agreement from the Johns Hopkins University School of Medicine and the Neurofibromatosis Therapeutic Acceleration Program (NTAP). Its contents are solely the responsibility of the authors and do not necessarily represent the official views of the Johns Hopkins University School of Medicine. The work has also been partially supported by the Spanish Ministry of Science and Innovation, Carlos III Health Institute (ISCIII) (PI20/00228) Plan Estatal de I+D+ I 2013–2016, co-financed by the FEDER program – a way to build Europe –; and by the Government of Catalonia (2017-SGR-496) and CERCA Program/Generalitat de Catalunya. MM-L has been supported by Fundación PROYECTO NEUROFIBROMATOSIS. We also would like to thank all members of the Lázaro lab and the IGTP core facilities and their staff for their contribution and technical support: Flow Cytometry (Gerard Requena and Marco A. Fernández); High Content Genomics and Bioinformatics Core Facility (Lauro Sumoy; Raquel Pluvionet); High Performance Computing. We are also grateful to all patients that donated neurofibromas for this project and to the Fundación Proyecto Neurofibromatosis, the Asociación de Afectados de Neurofibromatosis (AANF) and the Catalan Neurofibromatosis Association (ACNefi) for their constant support.

