## Supplemental material for "Unbalancing cAMP and Ras/MAPK pathways as a therapeutic strategy for cutaneous neurofibromas"

### List of supplemental Figures, Tables and Files

#### Supplemental Figures

**Supplemental Figure 1.** Setting up SC-FB co-culture conditions.

**Supplemental Figure 2.** p75 expression in co-cultures as a control of co-culture composition for RNA-seq analysis.

**Supplemental Figure 3.** Validation of the SC-FB crosstalk signature in an iPSC-derived neurofibromasphere model.

**Supplemental Figure 4.** Toxicity effect of single drugs tested on NF1 patient skin fibroblasts.

**Supplemental Figure 5.** Ogerin treatment increases cAMP levels in cNF-derived SCs.

**Supplemental Figure 6.** Toxicity effect of Selumetinib-Ogerin co-treatment in NF1 skin fibroblasts.

**Supplemental Figure 7.** Effect of Selumetinib and Ogerin treatments and co-treatments on cell death in primary SC cultures.

#### Supplemental Tables

**Supplemental Table 1.** List of candidate genes and activators and inhibitors used.

**Supplemental Table 2.** Summary of cNFs used in functional assays using single cultures of SCs, FBs and SC-FB co-cultures.

#### Supplemental Files (available upon request)

**Supplemental File S1**. Transcriptional signature specific to SC-FB interaction.

DEG de real vs virtual co-cultures

**Supplemental File S2**. Enrichment analysis using the SC-FB interaction gene signature

**Supplemental File S3**. Luminex measurements of supernatants in single cultures and co-cultures

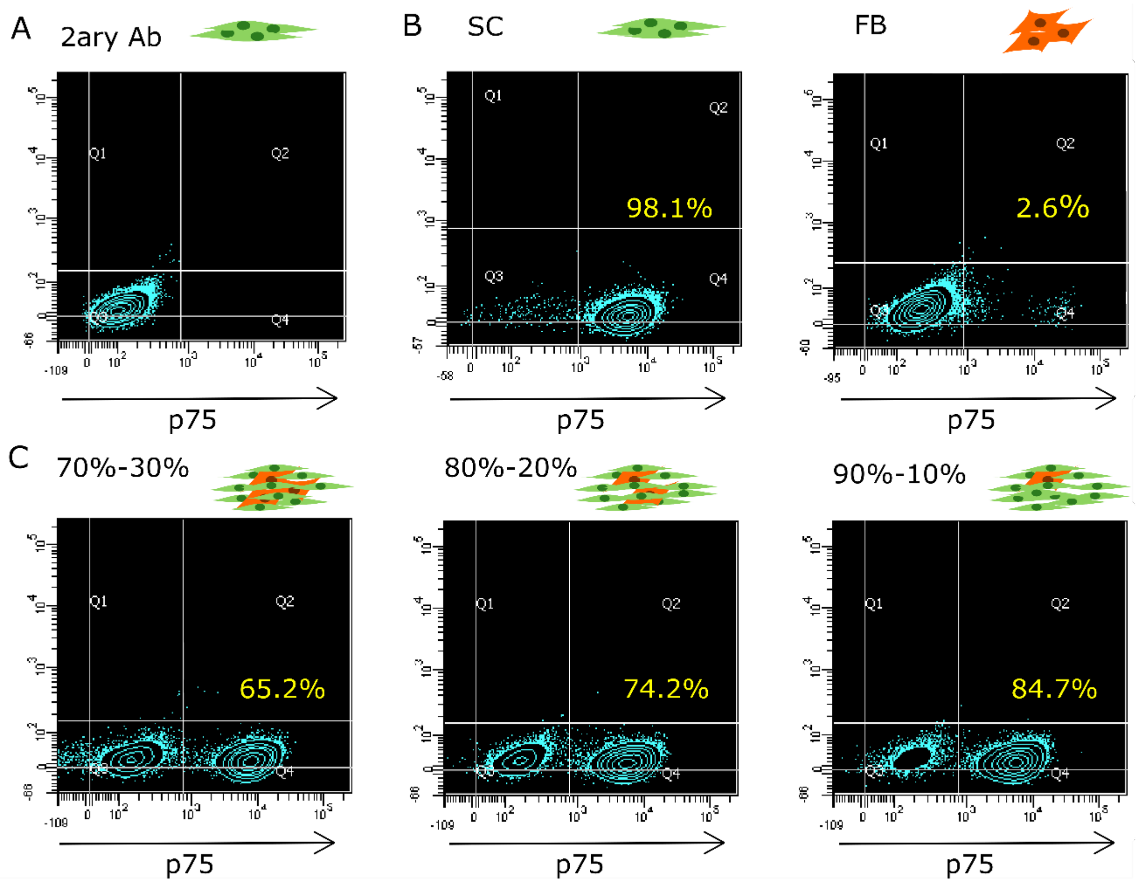


**Supplemental Figure 1. Setting up SC-FB co-culture conditions.** Seeding 70% SC– 30% FB co-culture is enough to obtain a percentage of cells at 72 hours, closely resembling cNF composition. Flow cytometry analysis of p75 in single SC and FB cultures and different proportion of SCs and FBs co-cultures.

**A)** Control SCs incubated with the secondary antibody. **B)** Single SC and FB cultures incubated with p75 primary antibody and the secondary antibody. **C)** Percentages of p75 positive cells in SC-FB co-cultures after 72h, at different initial seeding conditions (70% SCs– 30% FB; the 80% SCs– 20% FB and the 90% SCs – 10% FB; respectively). Percentages of p75 positive cells are indicated in yellow. SC: Schwann cell; FB: Fibroblast. **Related to Figure 1**.

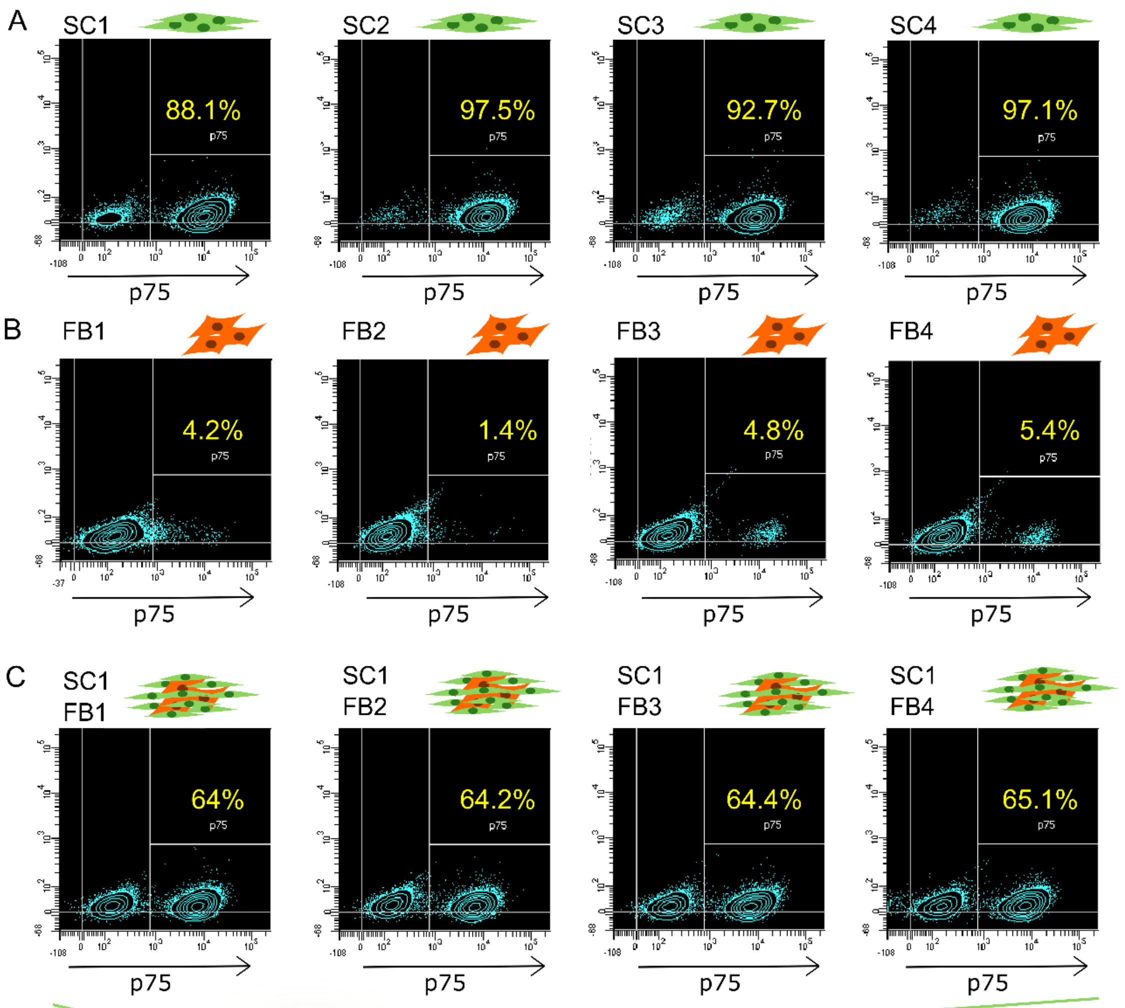


D

| | SC1 | SC2 | SC3 | SC4 |
|---|---|---|---|---|
| FB1 | 64 | 57.3 | 71.4 | 69.3 |
| FB2 | 64.2 | 53.8 | 68.2 | 66 |
| FB3 | 64.4 | 60.6 | 69 | 76.2 |
| FB4 | 65.1 | 59.9 | 69.6 | 69.6 |

**Supplemental Figure 2. p75 expression in co-cultures as a control of co-culture composition for RNA-seq analysis.** Flow cytometry analysis for p75 of single SC and FB cultures and SC-FB co-cultures of four single cNF-derived SC cultures, four single cNF-derived FB cultures, and four different co-culture combinations at 72 hours are shown. The exact percentage of cells expressing p75 is shown in yellow **A)** Four independent single SC cultures. **B)** Four independent single FB cultures. **C)** Four independent co-cultures combinations using SC1 Schwann cells and FB from the four independent cNFs (FB1, FB2, FB3, and FB4). **D)** Summary of the different percentages of p75-expressing cells in the sixteen different co-culture combinations at 72 hours. SC: Schwann Cells; FB: Fibroblasts. **Related to Figure 1 and 2**.

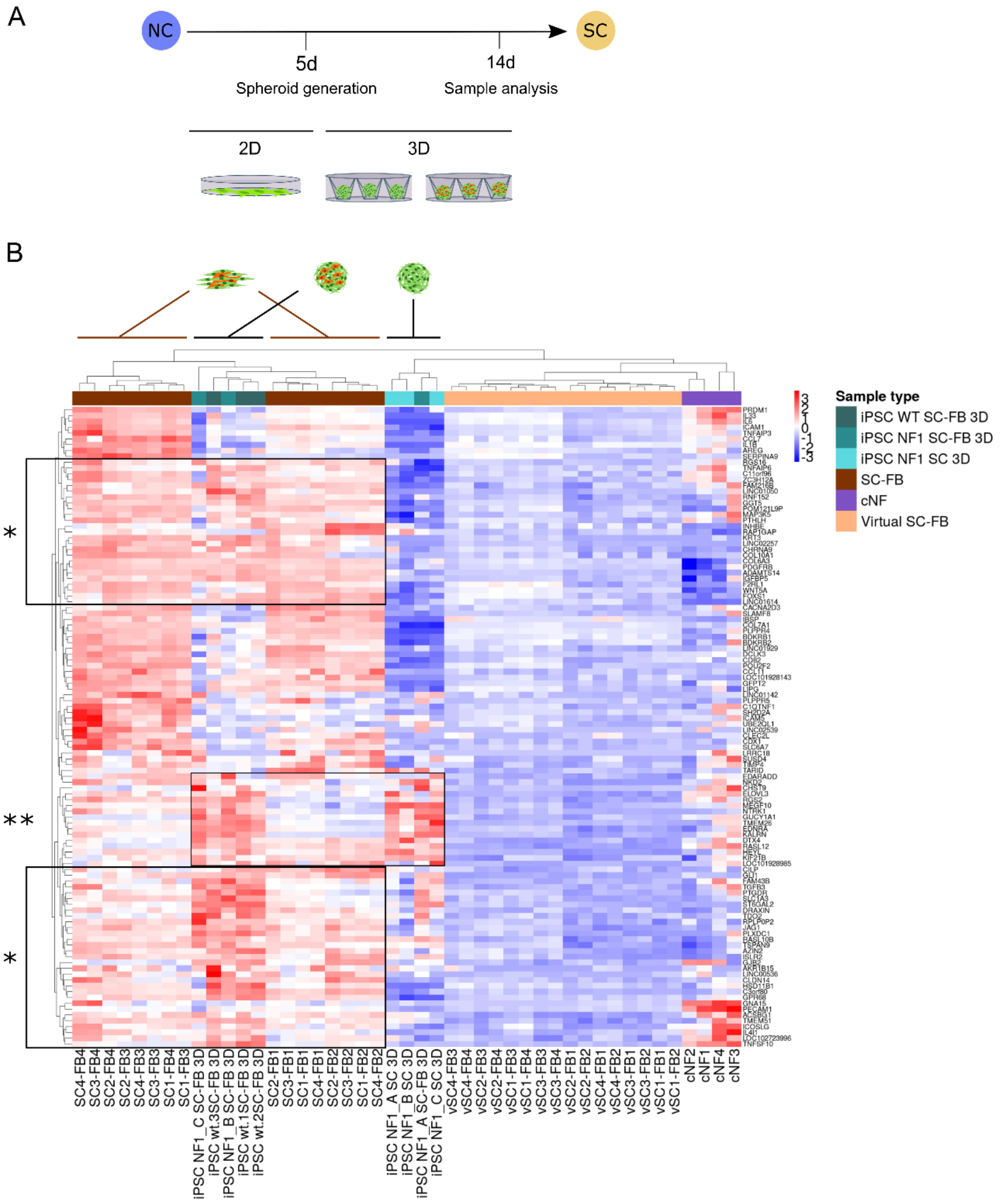


**Supplemental Figure 3. Validation of the SC-FB crosstalk signature in an iPSC-derived neurofibromasphere model.** iPSC-derived SC-FB 3D co-cultures express most genes upregulated in cNF-derived SC-FB co-cultures. **A)** Schematic representation of the iPSC-based SC or SC-FB model in 3D from Mazuelas et al. 2022. **B)** Heatmap showing the unsupervised cluster analysis of differentially upregulated genes in SC-FB co-cultures for different samples: iPSC-derived *NF1*(-/-) SC 3D culture (light turquoise); iPSC-derived SC-FB 3D co-culture (dark turquoise) spheroids generated from *NF1*(+/+) (WT, dark) and *NF1*(-/-) (NF1) iPSCs; real (brown) and virtual (salmon) SC-FB co-cultures; and in cNFs (purple). The expression color ranges from dark blue, showing down-regulated genes, to red, showing up-regulated genes. WT: wild type, SC: Schwann Cell, FB: Fibroblast; NC: Nerual Crest. * expression profile that occurs due to SC-FB interaction, wether in 2D o 3D. ; ** expression profile that occurs due to SC-SC interaction. **Related to Figure 2**.

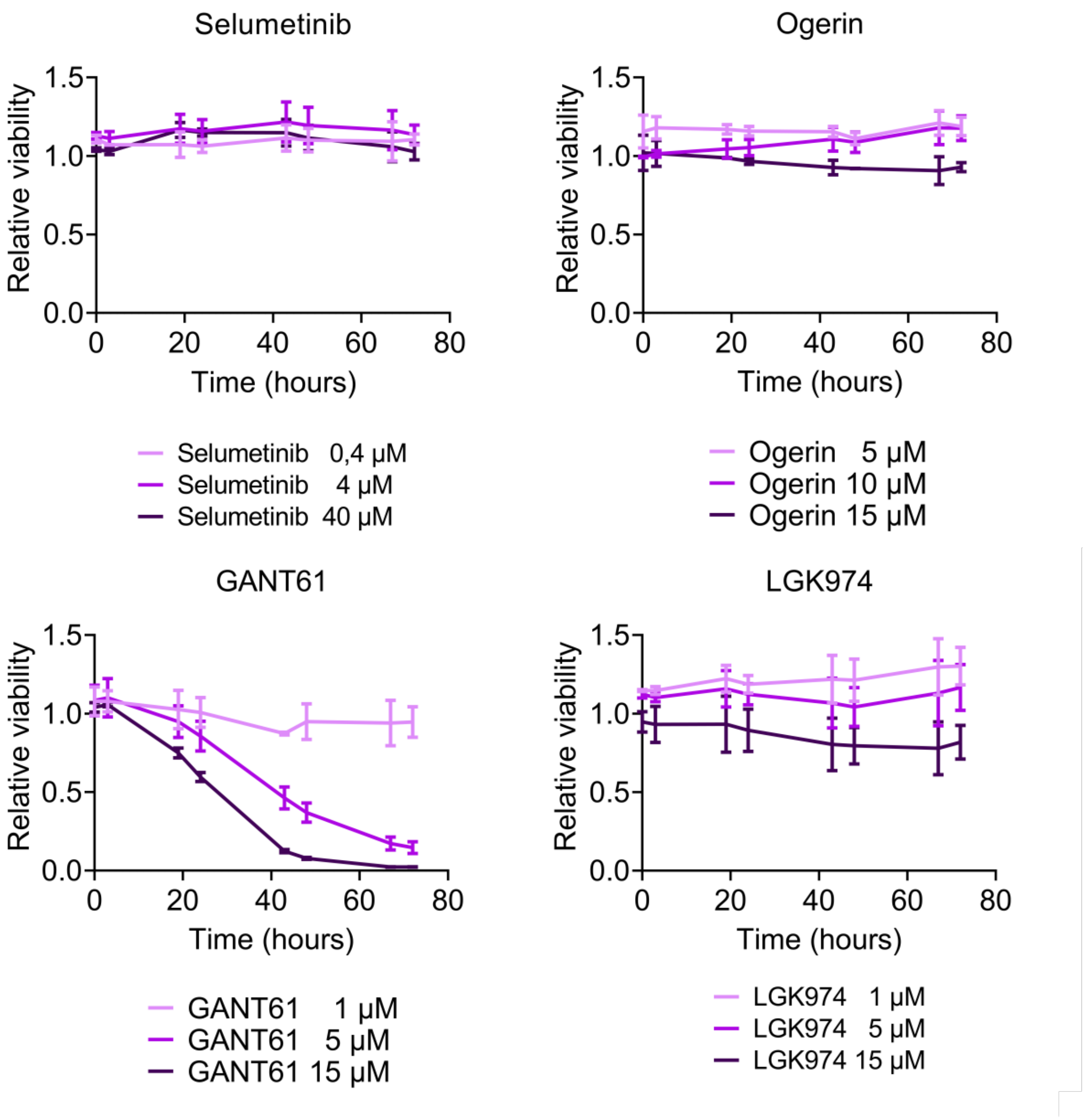


**Supplemental Figure 4. Toxicity effect of single drugs tested on NF1 patient skin fibroblasts.**

NF1 patient skin fibroblasts were plated and 24h later treated with different doses of Selumetinib, Ogerin, GANT61, and LGK974. Cell viability was monitored throughout 72h using RealTime-Glo MT Cell Viability Assay. Data are expressed as mean +/- SEM from three different fibroblast cultures, related to DMSO-treated control cells. **Related to Figure 5.**

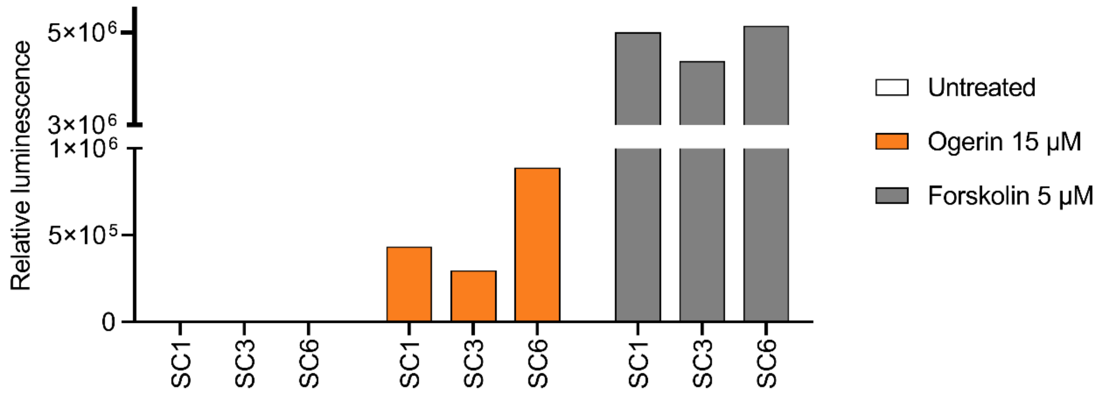


**Supplemental Figure 5. Ogerin treatment increases cAMP levels in cNF-derived SCs**. Intracellular cAMP levels were quantified using the cAMP-Glo Max Assay (Promega). SCs were plated in SCM without IBMX with Forskolin and maintained at 37ºC under a 10% CO2 atmosphere for 24 hours. At this point, cells were treated with induction buffer containing Ogerin or Forskolin or none for 30 minutes at room temperature. cAMP-Glo Max Assay was performed following manufacturer's instructions, and Luminescence was monitored on a Varioskan Flash plate reader (Thermo). Relative luminescence represents the values from treated compared to untreated. **Related to Figures 5 and 6.**

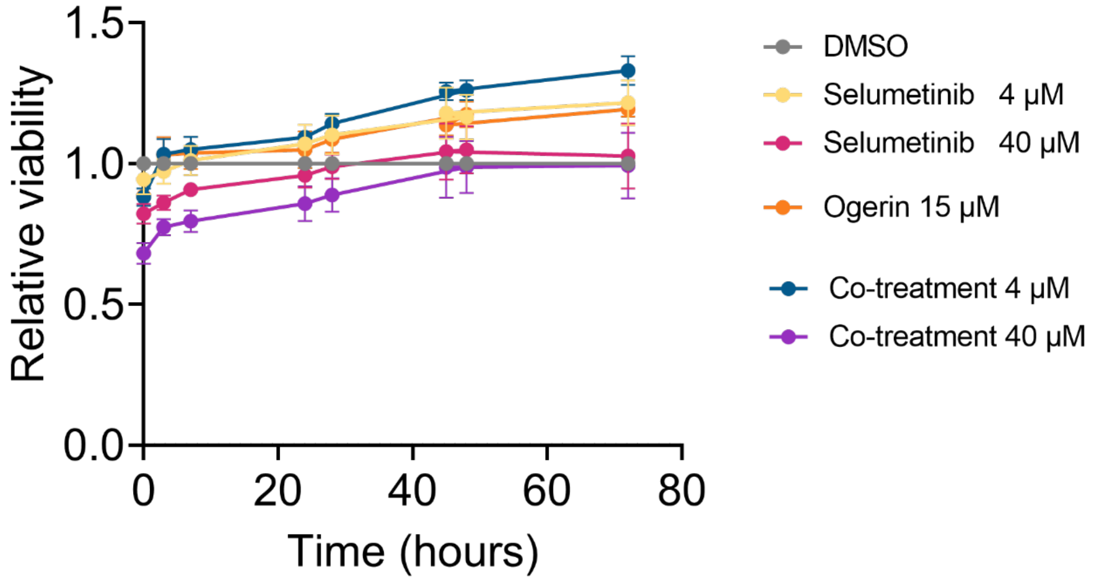


**Supplemental Figure 6. Toxicity effect of Selumetinib-Ogerin co-treatment in NF1 skin fibroblasts.** Cells were plated and 24h later treated with single drugs (4 µM Selumetinib, 40 µM Selumetinib, 15 µM Ogerin) or combination of drugs (4 µM Selumetinib and 15 µM Ogerin; 40 µM Selumetinib and 15 µM Ogerin). Cell viability was monitored throughout 72h using RealTime-Glo MT Cell Viability Assay. Data are expressed as mean +/- SEM from three different fibroblast cultures, related to DMSO-treated control cells. **Related to Figure 6.**

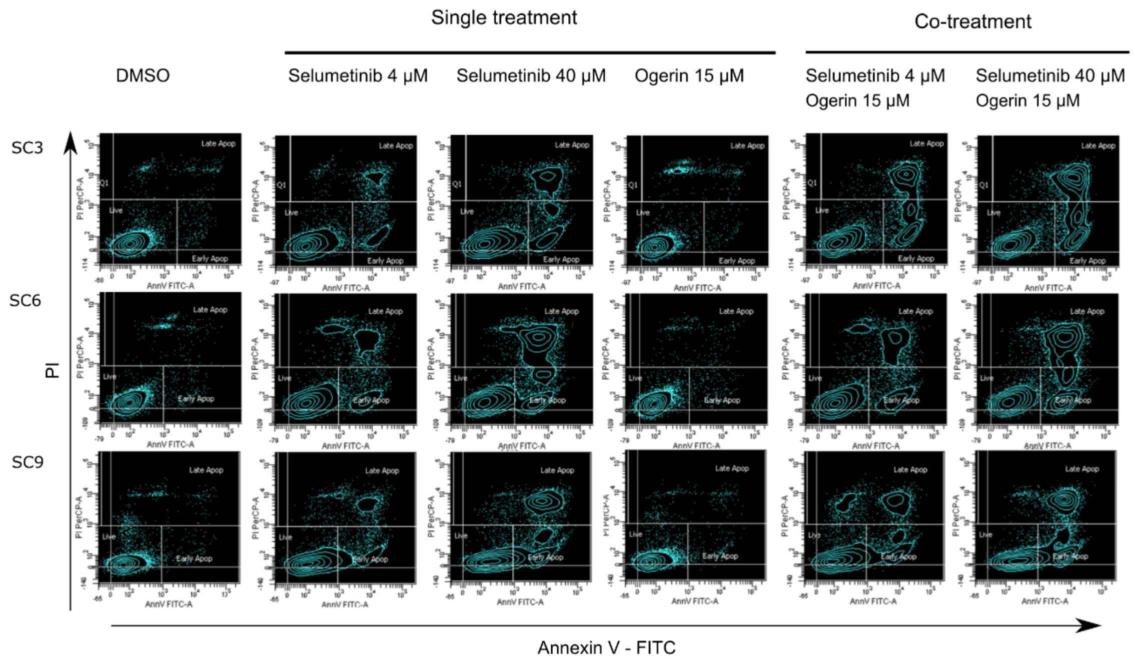


**Supplemental Figure 7. Effect of Selumetinib and Ogerin treatments and co-treatments on cell death in primary SC cultures.** Cytometry plots showing Annexin V and Propidium Iodide (PI) staining after 48h of treatments. Cells were plated and 24h later treated with vehicle (DMSO), single drugs (4 µM Selumetinib, 40 µM Selumetinib, 15 µM Ogerin) or co-treatment of drugs (4 µM Selumetinib and 15 µM Ogerin; 40 µM Selumetinib and 15 µM Ogerin) for 48 h. At this point, apoptosis was quantified by flow citometry using the Annexin V FITC Apoptosis detection kit (Invitrogen). **Related to Figure 6.**

**Supplemental Table 1. List of candidate genes and activators and inhibitors used.**

| Candidate | Signaling pathway | Inhibitors/ Activators | | |
|---|---|---|---|---|
| | | Type | Name | Reference |
| AREG | EGF TGFalpha | Neutralizing Antibody | Human Amphiregulin mAb (Clone 31221) | R&D MAB262-SP |
| TGFb3 | TGFb | Neutralizing Antibody | TGFb3 mAb (Clone 20724) | R&D MAB243-SP |
| JAG1 | Notch | Neutralizing Antibody | Human Jagged 1 polyclona Ab | R&D AF1277-SP |
| GLI1 | Sonic Hedgehog | Inhibitor | GANT61 | TOCRIS 3191 |
| WNT5A | Wnt | Inhibitor | LGK-974 | Selleckchem S7143 |
| GPR68 | AMPc | Activator | Ogerin | TOCRIS 5722 |
| TGFA | TGFalpha | Activator | Recombinant Human TGF alpha | Abcam ab233681 |
| Positive control | ERK | Inhibitor | Selumetinib | TOCRIS 6815 |

**Supplemental Table 2. Summary of cNFs used in functional assays using single cultures of SCs, FBs and SC-FB co-cultures.**

| | | | Real Time Cell Titer-Glo Viability Assay | | | Proliferation Assay (EdU) | Real Time Cell Titer-Glo Viability Assay | Apoptosis Assay | cAMP-Glo Max Assay |
|---|---|---|---|---|---|---|---|---|---|
| **NF1 patient ID** | **Sex** | **SC-Fb co-culture Fig2** | **SC Fig5C** | **FB Fig5C** | **SC-FB Fig5C** | **Fig5D** | **SC Co-treatment Fig6A** | **SC Co-treatment Fig6C** | **SC Ogerin treatment FigS6** |
| 1 | M | X | X | X | X | | | | X |
| 2 | M | X | | | | | | X | |
| 3 | F | X | X | X | X | X | X | X | X |
| 4 | F | X | | | | X | | | |
| 5 | F | | | | | | X | | |
| 6 | M | | | X | | | | X | X |
| 7 | M | | | | | | X | | |
| 8 | M | | X | | X | X | | | |
